# Dual RNAseq highlights the kinetics of skin microbiome and fish host responsiveness to bacterial infection

**DOI:** 10.1101/2020.08.26.247221

**Authors:** J. Le Luyer, Q. Schull, P. Auffret, P. lopez, M. Crusot, C. Belliard, C. Basset, Q. Carradec, J. Poulain, S. Planes, D. Saulnier

## Abstract

**Background:** *Tenacibaculum maritimum* is a worldwide-distributed fish pathogen known for causing dramatic damages on a broad range of wild and farmed marine fish populations. Recently sequenced genome of *T. maritimum* strain NCIMB 2154^T^ provided unprecedented information on the possible molecular mechanisms involved in virulence for this species. However, little is known on the dynamic on the infection *in vivo*, and information are lacking on both the intrinsic host response (gene expression) and its associated microbiome community. Here, we applied complementary omic approaches, including dual RNAseq and 16S rRNA gene metabarcoding sequencing using Nanopore and short-reads Illumina technologies to unravel the host-pathogens interplay in experimental infection system using the tropical fish *Platax orbicularis* as model.

**Results:** We show that *T. maritimum* transcriptomic landscape during infection is characterized by an enhancement of antibiotic catalytic and glucan catalytic functions while decreasing specific sulphate assimilation process, compared to *in vitro* cultures. Simultaneously, fish host display a large palette of immune effectors, notably involving innate response and triggering acute inflammatory response. In addition, results suggest that fish activate adaptive immune response visible through stimulation of T-helper cells, Th17, with congruent reduction of Th2 and T-regulatory cells. Fish were however largely sensitive to infection, and less than 25% of them survived after 96hpi. These surviving fish showed no evidence of stress (cortisol levels) as well as no significant difference in microbiome diversity compared to control at the same sampling time. The presence of *Tenacibaculum* in resistant fish skin and the total absence of any skin lesion suggest that these fish did not escape contact with the pathogen but rather prevent the pathogen entry. In these individuals we detected the up-regulation of specific immune-related genes differentiating resistant from control at 96hpi, which suggests a possible genomic basis of resistance while no genetic variations in coding regions was reported.

**Conclusion:** Here we refine the interplay between common fish pathogens and host immune response during experimental infection. We further highlight key actors of defense response, pathogenicity and possible genomic bases of resistance to *T. maritimum*.

## Background

Pathogens remain a significant threat to biodiversity, livestock farming and human health [1]. Host–pathogen interactions rely on a complex balance between host defenses and pathogen virulence. Through constant selective pressure, pathogens evolve mechanisms in order to overcome the host immune system, likewise the host adapt to counteract and limit pathogen virulence. Although changes in gene expression as a result of host–pathogen interactions appear to be common [2–4], the mechanisms involved often remain poorly understood. A more in-depth understanding of host–pathogen interactions has the potential to improve our mechanistic understanding of pathogenicity and virulence, thereby defining novel preventive, therapeutic and vaccine targets [5].

Dual RNAseq sequencing strategy fills the need of simultaneously assess the host and the pathogens genes expression [6–8]. Studies applying these approaches in fish bacterial infection systems have bloomed recently and show promising in deciphering the complexity of host-pathogen interplay [9–11] yet, these studies did not simultaneously explore dysbiosis and associated changes of microbiome communities. In nature, co-occurrence of multiple pathogen species (co-infection) is frequent. Species interactions might be neutral, antagonistic or facilitative and most often shape strain virulence plasticity resulting in increased disease virulence [12–14]. Despite its commonness, remarkably few studies have explored such models, *i*.*e*. when host interact simultaneously with multiple pathogens co-infection [15]. During tenacibaculosis outbreak in platax, *T. maritimum* burden is also commonly associated with other pathogen co-occurrences, namely *Vibrio* spp [16]. Nevertheless, such approach is dramatically impaired by the unbalanced representation of each compartment sequences, most often favouring the host compartment [8,17]. This bias can be minimized by specific library preparation (*i*.*e*. mRNA depletion), *in silico* normalization procedures, and/or by investigating models where pathogens burden is high.

*Tenacibaculum maritimum* is a worldwide-distributed fish pathogen, known for its lethal consequences on a broad range of wild and farmed marine fish populations. Major efforts have been undertaken to lessen the effect of the pathogen and/or increase fish immune resistance [18], yet the mechanisms of the infection and the response of fish remain largely unknown which significantly hold back development of aquaculture sectors. Nevertheless, recent sequencing of *T. maritimum* strain NCIMB 2154^T^ genome provided unprecedented information on the putative molecular mechanisms involved in virulence [19]. Authors note for instance that *T. maritimum* display a large array of evolutionary conserved stress resistance related effectors as well as expanded capacity of iron mobilisation [19].

*Tenacibaculum maritimum* adheres and rapidly colonizes mucosal surfaces [16,20]. Infected fish show multiplication of *T. maritimum* on their external tissues leading to severe skin lesions and following fish rapid death [16]. Therefore, as for other infection systems, the mucosal surfaces, here mainly skin mucus, is considered as the first barrier against pathogens [21]. This physical and chemical barrier constituted by mucus also includes the presence of host immune effectors (innate and adaptive) that orchestrate a complex interaction network between with the commensal bacterial community [22,23]. The identification of these multi-specific interactions within the mucus brings to the fore the microbiome as the cornerstone of host-pathogen interactions [21,24]. Indeed, dysbiosis (i.e., the imbalance or alteration of the microbial ecosystem leading to a diseased status) is directly involved in the severity of a disease [25–27]. Recent studies on zebrafish (*Danio rerio*) raised in axenic conditions or in the presence of probiotic bacteria underlined the crucial role of the microbiota on the development of the immune system, mucosal homeostasis and resistance to stress and pathogens ([28], for review see [29]).

In French Polynesia, recurrent tenacibaculosis infections have been the major obstacle to local fish production sustainability. Indeed, *T. maritimum* affects the only locally farmed fish, the Orbicular batfish (*Platax orbicularis*) leading to very high mortality rates shortly after transferring hatchery fingerlings to off-shore marine cages. Here we combined dual RNAseq and 16S rRNA metabarcoding sequencing approaches to investigate the molecular responses of the host and the microbiome communities (genes expression and microbiome composition) simultaneously during the infection and the recovery phases to *T. maritimum* using the orbicular batfish as a model. We also integrated comparison to *in vitro* liquid cultures of *T. maritimum* to refine our knowledge of virulence-related genes and explored genomic and genetic bases of resistance in *P. orbicularis*.

## Methods

### a) Animal husbandry

Platax fingerlings used in the bacterial challenge were 58 days old (days post-hatching; dph). Fish were obtained from a mass tank spawning of 6 females and 8 males induced by desalinisation. Broodstock include wild individuals caught in French Polynesia that has been maintained at the Centre Ifremer du Pacifique (CIP) hatchery facility for seven years, under the supervision of the direction des ressources marines (DRM). Eggs were randomly distributed into six black circular fiberglass tanks of 210 L with 50 eggs.L^−1^ in order to achieve an average density of 30 larvae.L^−1^. Half of the tank were then reared in conditions that followed standard procedures implemented in the CIP facility (open water system, normal salinity around 36 psu), named “Standard” [18]. The other half was reared in conditions that were supposed to be optimal according to previous experiments, named “Recirculated”. Indeed, animals were bred in a recirculating system which was desalinated to 24 psu until day 34 where salinity was progressively raised to normal (36 psu). Moreover, commercial clay (Clay Bacter ®) was added daily at a rate of 1g/d/m^3^ per percent of hourly water renewal from day 1 to day 19, the beginning of living prey weaning period. In addition, an input of probiotic *Pseudoalteromonas piscicida B1* (local strain), produced in CIP facilities (see supplementary methods), was realised daily in fish fed (0.5mL/day/tank) and in the water (0.5 ml/day/tank) at a concentration of 10^9^cfu/ml of bacterial suspension from day 1 to day 57. At the end of the larval phase, day 20, 700 fingerlings/tank were randomly kept. When they reached an average weight of 1g, they were sorted to get rid of the queue and head batch and kept at a density of 200/tank (1g/L). It is however relevant mentioning that around day 40, fish started to show a decrease of appetite and heavy mucus losses even if there was no mortality, mainly in the recirculating system. After an analysis of water flora, it was shown that *Vibrio harveyi* was present but no *T. maritimum*.

Platax larvae were fed 4 times a day with living preys (*Brachonius sp*. and *Artemia spp*.) before being weaned from day 16 to 23. Fingerlings were then fed with commercial micro-pellets ranging from 0,3 to 1 mm for the range Micro-Gemma and Gemma (Skretting, Stavanger, Norway) and 1-1.3 mm for Ridley (Le Gouessant, Lamballe, France) according to the standard previously established [18]. Seawater provided to both systems was pumped from the lagoon, filtered with 300 µm sand filter and two 25 and 10 µm mesh filters and UV treated (300 mJ/cm^2^). Recirculating system included a 500L biological filter to regulate level of ammonia and nitrite. All tanks were supplied with saltwater held at 28,4 ± 0.3°C at constant photoperiod (12L: 12D) and oxygen saturation was maintained above 60% in the tanks with air distributed via air stone. Water renewal ranged from 36 to 360 L/h and new water input in the recirculating system was of 11 ± 1 %. Levels of ammonia and nitrite were monitored once a week by spectrophotometry (HANNA Instruments ®) to assess bio filter performance. Temperature, salinity and dissolved oxygen were measured daily (YSI ®) and the unfed and fecal materials were removed once a day.

### b) Bacterial challenge

We used strain TF4 for the experimental infection on 58 dph fingerlings. TFA4 strain was isolated from the skin of an infected *Platax orbicularis* in French Polynesia in 2013 and was shown to belong to *Tenacibaculum maritimum* by whole-genome sequencing, displaying an average nucleotide identity of 99.6 % with the reference strain NCIMB 2154^T^ [30]. TFA4 strain was cultivated in nutrient Zobell medium (4 g L^−1^ peptone and 1g L^−1^ yeast extract Becton, Dickinson and Company, Sparks, MD in filtered and UV-treated sea water) under constant agitation (200 rpm) at 27°C for 48h. On the infection day, 50 juvenile *Platax orbicularis* were transferred into 40 L-tanks supplied with air and infected by an addition of 10 mL of bacterial suspension of TFA4 strain in the tank water. Final bacterial concentration in the 40 L-tanks tanks, determined by “plate-counting” method, reached 4.10^4^ CFU.mL^−1^. Moreover, for each rearing condition, 16 to 17 fish were randomly selected from each tank to be transferred in one 40L-tank to form mock-treated group, referred as control (N = 50 fish) and an addition of 10 mL of Zobell medium was done. After 2 hours of bathing, fish were caught with a net, rinsed by successive passage in two buckets of 40L filled with clean filtered UV-treated seawater, before being transferred back into their respective tanks. Twice a day, one third of the water was changed with filtered UV-treated seawater to maintain good water quality and be able to inactivate *T. maritimum* in sewage by bleach treatment. Moreover, at those times, dead animals were collected and recorded. After 115 hours post-infection (hpi), all infected animals were considered as survivors and the challenge ended. All the remaining fish were euthanized.

### c) Animal sampling

We randomly sampled five individuals per tank at 58 dph (average weight of 7.21g ± 0.28 se). Sampling used for the experiment consisted in five individuals per tank at 24hpi and 96hpi (Figure 1A). To increase our sampling dataset for controls we included “standard” control group (see previous section, one tank) together with “recirculated” condition (one tank) individuals. Consequently, our design consisted in four groups, namely *control*_*24h*_ (N=10 individuals, replicates tanks), *control*_*96h*_ (N=10 individuals, replicates tanks), *infected*_*24h*_ (N=15 individuals, triplicates tanks) and *resistant*_*96h*_ (N=15 individuals, triplicates tanks). For each sampling, at 24hpi and 96hpi, individuals were lethally anaesthetized using benzocaine bath (150 mg.L^−1^) and a lateral photograph was taken using a digital fixed camera (Leica Microsystems; Figure 1B and C). Microbiome and host sampling consisted in gentle fish skin smears with sterile swabs. Swabs were directly placed in TRIZOL Reagent (Life Technologies) on ice to prevent RNA degradation. Swabs were disrupted using a mixer mill MM200 (Retsch) for 5 min at a frequency of 30 Hz and stocked at −80°C for latter analysis. In parallel, water was also sampled in each tank but was not included in the analysis due to very low DNA yield.

**Figure 1:**
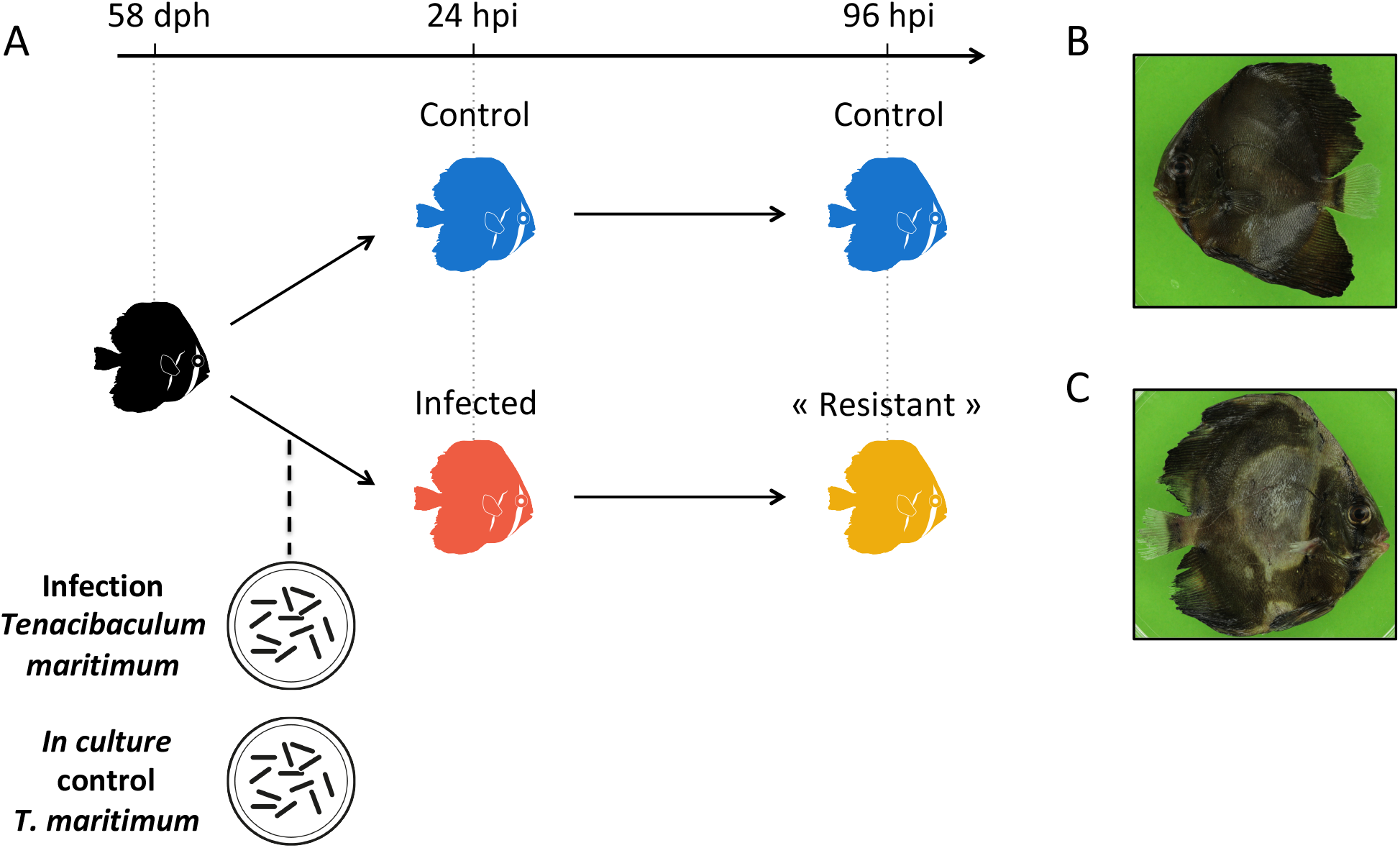
Experimental design and individuals’ photographs. **A)** Experimental infection was conducted at 58 dph, after random sampling of five individuals per tank to assess initial weight. At 24 and 96 hours hpi, five individuals per tank (N= 15 individuals, *infected*_*24h*_; N = 15 individuals, *resistant*_*96h*_; N = 10 individuals, *control*_*24h*_ and N = 10 individuals, *control*_*96h*_) were sampled by swabs. Same individuals serve for host and microbiome transcriptomic and for microbiome metabarcoding. B) Photograph of control fish (control_*24h*_); C) Photographs of infected fish (infected_*24h*_) showing typical skin lesions associated with tenacibaculosis.

### d) *T. maritimum in vitro* liquid culture sampling

The TFA4 strain was cultivated in 6 mL of Zobell medium under constant agitation (200 rpm) at 27°C for 48h, following exact same procedure and time of incubation than the culture used for bacterial challenge. Five culture replicates were performed. For each replicate, 4 mL at 10^8^ CFU/mL were centrifuged 5 min. at 10,000g and at room temperature. Three inox beads and 2 mL of TRIZOL (Life technologies) were quickly added to each bacterial pellet and cells were immediately disrupted using a mixer mill MM200 (Retsch) for 5 min. at a frequency of 30 Hz to prevent RNA degradation. RNA was extracted following manufacturer’ instructions, using a high salt precipitation procedure (0.8 M sodium citrate and 1.2 M NaCl per 1 ml of TRIZOL reagent used for the homogenization) in order to reduce proteoglycan and polysaccharide contaminations. Quantity, integrity and purity of total RNA were validated by both Nanodrop readings (NanoDrop Technologies Inc.) and BioAnalyzer 2100 (Agilent Technologies). DNA contaminants were removed using a DNAse RNase-free kit (Ambion). A total of five RNA samples (1.048 ± 0.019 µg) were further dried in RNA-stable solution (ThermoFisher Scientific) following manufacturer’s recommendations and shipped at room temperature to McGill sequencing platform services (Montreal, Canada). One library was removed prior sequencing because it did not meet the minimal quality requirements.

### e) Fish mortality and cortisol measurements

Mortality was recorded at 0, 19, 24, 43, 48, 67, 72, 91, 96 and 115hpi. We used the non-parametric Kaplan-Meier approach for estimating log-rank values implemented in the survival R package [31]. Differences in survival probability was considered significant for *P* < 0.05. We assessed stress levels in fish by measuring scale cortisol content [32]. The scales of both flanks of each individual were collected, and subsequently washed and vortexed three times (2.5 min; 96% isopropanol) in order to remove external cortisol that takes its source in mucus. Residual solvent traces were evaporated under nitrogen flux and samples frozen at −80°C. To ensure scales were dry, they were lyophilized for 12 hours before being grounded to a powder using a ball mill (MM400, Retsch GmbH, Germany). Cortisol content was extracted from ∼50 mg of dry scale powder by incubation in 1.5mL of methanol (MeOH) on a 30°C rocking shaker during 18 hr. After centrifugation at 9500g for 10 min, the supernatant was evaporated a rotary evaporator and reconstituted with 0.2 mL of EIA buffer provided by the Cortisol assay kit (Neogen® Corporation Europe, Ayr, UK). Cortisol concentrations were determined in 50 µL of extracted cortisol by using a competitive EIA kits (Neogen® Corporation Europe, Ayr, UK) according to previously published protocol [33]. Differences across groups were tested by a two-way ANOVA and following Tukey’s HSD post-hoc after validation of normality and homoscedasticity. Differences were considered significant when *P* < 0.05.

### f) RNA and DNA extraction and sequencing

#### Dual RNAseq

Total RNA was extracted using the same procedure than described above. RNA was then dried in RNA-stable solution (ThermoFisher Scientific) following manufacturer’s recommendations and shipped at room temperature to McGill sequencing platform services (Montreal, Canada). Ribo-Zero rRNA removal kit (Illumina, San 260 Diego, Ca, USA) was used to prepare mRNA, rRNA-depleted, libraries that were multiplexed (13-14 samples by lane) and sequenced on HiSeq4000 100-bp paired-end (PE) sequencing device. Infected individuals 24hpi were sequenced twice for insuring sufficient coverage (Table S1).

#### Short-reads 16S rRNA MiSeq microbiome sequencing

Total DNA was extracted from the same TRIZOL Reagent (Life Technologies) mix than described above. DNA quantity/integrity and purity were validated using both a Nanodrop (NanoDrop Technologies Inc.) and a BioAnalyzer 2100 (Agilent Technologies). The V4 region was amplified by PCR using modified 515F/806rb primers constructs (515F: 5’-GTGYCAGCMGCCGCGGTAA-3’; 806rb: 5’-GGACTACNVGGGTWTCTAAT-3’), recommended for microbial survey [34]. Amplicons libraries were multiplexed and sequenced on a single lane of MiSeq 250bp PE Illumina machine at Genome Québec McGill, Canada. Details of sequencing statistics are provided in Table S2.

#### Full 16S rRNA Nanopore sequencing

For a broad range amplification of the 16S rRNA gene, DNA was amplified using the 27F/1492R barcoded primers products (27F: 5’-AGAGTTTGATCMTGGCTCAG-3’; 1492R: 5’-TACGGYTACCTTGTTACGACTT-3’). We included in the PCR experiment eight randomly selected individuals from *infected*_*24h*_ group, two negative PCR controls (clean water) and one positive control (Acinetobacter DNA).

The PCR mixtures (25 μl final volume) contain 10 ng of total DNA template or 10 µl of water, with 0.4 μM final concentration of each primer, 3% of DMSO and 1X Phusion Master Mix (ThermoFisher Scientific, Waltham, MA, USA). PCR amplifications (98 °C for 2 min; 30 cycles of 30 s at 98 °C, 30 s at 55 °C, 1 min at 72 °C; and 72 °C for 10 min) of all samples were carried out in triplicate in order to smooth the intra-sample variance. Triplicates of PCR products were pooled and purified by 1x AMPure XP beads (Beckmann Coulter Genomics) cleanup. Amplicon lengths were measured on an Agilent Bioanalyzer using the DNA High Sensitivity LabChip kit then quantified with a Qubit Fluorometer.

An equimolar pool of purified PCR products (excepted for negative controls) was done and one sequencing library was finally prepared from 100 ng of the pool using the 1D Native barcoding genomic DNA protocol (with EXP-NBD103 and SQK-LSK108) for R7.9 flow cells run (FLO-MAP107) then sequenced on the MinION device. Details of sequencing statistics are provided in Supplementary Material.

### g) Microbiome communities analyses

#### Microbiome dynamics with MiSeq short-reads dataset

Raw reads were filtered to remove Illumina’s adaptors as well as for quality and length using Trimmomatic v.0.36 [35] with minimum length, trailing, and leading quality parameters set to 100 bp, 20, and 20, respectively. Remaining reads were analysed with functions implemented in QIIME2 platform v2019.10. Briefly, we used DADA2 algorithm [36] to cluster sequences in amplicon sequence variants (ASVs). The following ASVs were mapped against GreenGenes v13.9 99% OTUs database [37]. We explored alpha-diversity [Shannon, Fisher and Faith’s phylogenetic diversity (PD) indexes] and beta-diversity (Bray-Curtis, unweighted and weighted Unifrac distances) using phyloseq R package [38]. Dissimilarity between samples was assessed by principal coordinates analysis (PCoA). Differences in alpha-diversity were tested using pairwise Wilcoxon rank test and were considered significant when *P* < 0.01. Differences in beta-diversity were tested using PERMANOVA (999 permutations) as implemented in the adonis function of the vegan R package [39] and were considered significant when *P* < 0.01. We also searched for « core » microbiome in fish skin and considered as member of core microbiome ASVs that were present in all the individuals across all condition (infected, control and resistant). We finally searched for significant differences in specific ASV abundance across groups using Wald tests implemented in the DESeq2 R package [40]. We used ‘*apeglm*’ method for Log2FC shrinkage to account for dispersion and variation of effect size across individuals and conditions, respectively [41]. Differences were considered significant when FDR < 0.01 and |FC| > 2.

#### Microbiome diversity analysis with the Nanopore dataset

Sequences were called during the MinION run with the MinKnow software (v. 1.7.14). The demultiplexing and adaptor trimming were done with porechop tool (https://github.com/rrwick/Porechop) with the option discard_middle. For each barcode, all nanopore reads were mapped on the GreenGenes database (v.13.5, http://greengenes.lbl.gov) with minimap2 (v2.0-r191) with the pre-set options “map-ont” [42]. All reference sequences of the GreenGenes database covered by more than 0.01% of all read were kept for the next step. A second round of mapping (same parameters) was done on the selected references in order to aggregate reads potentially mis-assigned during the first round of mapping. SAMtools and BCFtools were used to reconstruct consensus sequences for each reference sequence covered with more than 10 nanopore reads with the following programs and options: mpileup -B -a -Q 0 –u; bcftools call -c --ploidy 1;vcfutils.pl vcf2fastq. Individuals and consensus sequences were blast against NCBI nt database (e-value < 10^−5^).

### h) Compartment specific differential expression analyses

#### Reads pre-processing

For each individual, raw reads were filtered using Trimmomatic v0.36 [43], with minimum length (60bp), trailing and leading (20 and 20; respectively). Filtered PE reads were mapped against a combined reference including the host’s transcriptome (See supplementary material for details of the transcriptome assembly; Table S1 and S3) and the genomes of *Alteromonas mediterranea* strain: AltDE1 (Genbank accession: GCA_000310085.1), and *Pseudoalteromonas phenolica* strain: KCTC 12086 (Genbank accession: GCA_001444405.1), *Tenacibaculum maritimum* strain: NCIMB 2154T (Genbank accession: GCA_900119795.1), *Sphingobium yanoikuyae* strain ATCC 51230 (Genbank accession: GCA_000315525.1), *Vibrio alginolyticus* strain: ATCC 17749 (Genbank accession: GCA_000354175.2,) and *Vibrio harveyi* strain: ATCC 43516 (Genbank accession: GCA_001558435.2). To prevent multi-mapping biases we used GSNAP v2017-03-17 [44] with minimum coverage fixed at 0.9, maximum mismatches allowed of 5 and removing non-properly paired and non-uniquely mapped reads (option “concordant_uniq”). Low mapping quality (MAPQ) were further removed using Samtools v1.4.1 [45] with minimum MAPQ threshold fixed at 5. A matrix of raw counts was built using HTSeq-count v0.9.1 [46]. Transcripts from host and bacteria species origin were then separated in different contingency tables using homemade scripts.

#### Host transcriptome analysis

Low coverage transcripts with count per million (CPM) < 1 in at least 9 individuals were removed, resulting in a total of 22,390 transcripts. Similarly, transcripts over-representation was assessed using *‘majSequences*.*R’* implemented in SARTools suite [47]. We used distance-based redundant discriminant analysis (db-RDA) to document genetic variation among groups and correlation with condition (infected or control), weight and time (24 and 96hpi) as the explanatory variables. Briefly, we computed Euclidean’s distances and PCoA using ‘*daisy*’ and ‘*pcoa*’ functions, respectively, implemented in the ape R package [48]. PCo factors (n = 6) were selected based on a broken-stick approach [49,50] and used to produce a db-RDA. Partial db-RDAs were used to assess the factor effect, controlling for the other factor variables. We tested the models and individual factors significance using 999 permutations. The effect was considered significant when *P* < 0.01.

Differential expression was assessed using the DESeq2 R package [40] using pairwise comparisons with Wald test. Logarithmic fold change (logFC) were shrinked using ‘*apeglm*’ method, implemented in DESeq2 R package [40], to account for dispersion and effect size across individuals and conditions [41]. Differences were considered significant when FDR < 0.01 and FC > 2. Group comparisons included *infected*_*24h*_ *vs control*_*24h*_ and *resistant*_*96h*_ *vs control*_*96h*_. Gene ontology (GO) enrichment was tested using GOAtools v0.6.5 [51] and the go-basic.obo database (release 2017-04-14) using Fisher’s test. Our background list included the ensemble of genes in the host transcriptome. Only GO terms with Bonferroni adjusted *P* < 0.01 and including at least three differentially expressed genes were considered. Significant GO enriched terms were used for semantic-based clustering in REVIGO (http://revigo.irb.hr/).

#### *Tenacibaculum maritimum* gene expression *in vitro* or during infection

A validation step for searching for transcripts over-representation was assessed using *‘majSequences*.*R’* implemented in SARTools suite [47], similarly to the fish transcriptome. Most represented sequences were attributed to *ssrA* coding genes, but represents less than 8% of the total library. We applied similar shrinkage method and pairwise comparisons (infected vs. *in vitro*). We used more stringent thresholds for the *T. maritimum* than for the host, as commonly observed in similar studies [8], and considered significant differences when FDR < 0.01 and FC > 4. Gene ontology (GO) enrichment was similar to the methods used for the host.

### i) Species specific weighted co-network gene expression analyses

We built a series of signed weighted co-expression network for the host and the bacteria compartments (using only for *infected*_*24h*_ individuals for including *V. harveyi, T. maritimum* and *P. phenolica*, independently) to cluster co-expressed genes and identify putative driver genes using WGCNA R package [52]. Variation in normalized counts were prior controlled using ‘*vst*’ method implemented in DESeq2 R package [40].

#### Host WGCNA analysis

We reduced the expression noise in the dataset by retaining only transcripts with minimum overall variance (> 5%). Briefly, we fixed a soft threshold power of 14 using the scale-free topology criterion to reach a model fit (|R|) of 0.80. The modules were defined using the ‘*cutreeDynamic*’ function (minimum 30 genes by module and default cutting-height = 0.99) based on the topological overlap matrix, a module Eigengene distance threshold of 0.25 was used to merge highly similar modules. For each module we defined the module membership (kME, correlation between module Eigengene value and gene expression values). We looked for significant correlation (Pearson’s correlation; *P* < 0.001) modules against physiological data including cortisol levels (pg.mg^−1^ of scales), fish weight (g) and condition (coded “1” for *control*_*24h*_, *control*_*96h*_ and *resistant*_*96h*_ and “2” for *infected*_*24h*_ condition, respectively). Gene ontology (GO) enrichment for each module was tested using same protocol and parameters than described above.

#### *T. maritimum* WGCNA analyses

Briefly, we conducted species-specific weighted co-expression network analyses and used bacterial species and host module eigenvalue to correlate genes modules. We fixed a soft threshold power of 20 using the scale-free topology criterion to reach a model fit (|R|) of 0.85. The modules were defined using the ‘*cutreeDynamic*’ function (minimum 30 genes by module and default cutting-height = 0.99) based on the topological overlap matrix, a module Eigengene distance threshold of 0.3 was used to merge highly similar modules. For each module we defined the module membership (kME, correlation between module Eigengene value and gene expression values). Individual module Eigengene values were computed using the *‘moduleEigengenes’* and used as metadata for further downstream correlation analyses. We finally computed a correlation matrix and focused on genes showing significant correlation (*P* < 0.05) to host main modules eigenvalues, namely module_host-turquoise_ and module_host-blue_. Gene ontology (GO) enrichment for each list of genes was tested using same protocol and parameters than described above.

### j) The genetic bases of fish resistance

We further explored the putative genetic variation between resistant and infected fish. We choose to focus on resistant fish because of their established phenotype, (*i*.*e*. survivor with no sign of lesions after bacterial challenge). We followed GATK recommendations for SNPs identification based on RNAseq data. Briefly, BAM files were pre-treated using ‘*CleanSam*’ function, duplicates were notified with the ‘*MarkDuplicates*’ function, and cigar string splitted with ‘*SplitNCigarReads*’ function. All functions are implemented in GATK v4.0.3.0 software [53,54]. Final SNPs calling was conducted with Freebayes v1.1.0 (https://github.com/ekg/freebayes) requiring minimum coverage of 15 and minimum mapping quality of 20, forcing ploidy at 2 and removing indels (‘*--no-indels’*) and complex polymorphisms (‘*--no-complex’*). The raw VCF file was filtered for minimum allele frequency (‘*--min_maf=0*.*2*’), minimum coverage (‘*--minDP=20’*) and allowing no missing data using Vcftools v0.1.14 [55]. We computed relatedness (‘*—relatedness2’*) within and among groups with Vcftools v0.1.14 [55]. We further used distance-based redundant discriminant analysis (db-RDA) to document genetic variation among groups and correlation with cortisol and condition and weight as the explanatory variables. Briefly, we computed Euclidean’s distances and PCoA using ‘*daisy*’ and ‘*pcoa*’ functions, respectively, implemented in the ape R package [48]. PCo factors (n = 6) were selected based on a broken-stick approach [49,50] and used to produce a db-RDA. We tested the model significance using 999 permutations, effect was considered significant when *P* < 0.01.

## Results

We compared infected (with *T. maritimum* TF4 strain) to mock-treated *P. orbicularis* groups (thereafter referred as control) in experimental infection conditions and individuals were sampled at 24 and 96 hpi. Surviving individuals in infected groups at 96hpi are referred as ‘resistant’ fish. Sampling consisted in individual skin swab for later DNA and RNA extraction. DNA served to assess microbiome communities (using 16S rRNA and nanopore technologies) while RNA served for genes expression analysis of the microbiome and the fish simultaneously (dual RNAseq). In addition, we compared genes expression of *T*.*maritimum* during the infection in vivo to in vitro liquid culture to detect putative virulence factors. Fish condition, survival and cortisol levels were also monitored.

### a) Fish weight, cortisol levels and mortality

Mortality rate in challenged fish reached 77.36 ± 18.35 (standard error; se) while no mortality was observed in control group (Kaplan-Meier analysis, *P* < 0.001; Figure 2A). Mortality started at 24hpi in infected group and no novel mortality even was observed after 72hpi. Cortisol levels in fish scales significantly vary across groups (ANOVA; *F* = 9.46; *P* < 0.01; Figure 2B). Overall cortisol levels were higher in the *infected*_*24h*_ group compared to all others groups (Tukey ‘s HSD; *P* = 0.01). Cortisol levels in *control*_*24h*_ group were also higher than both *control*_*96h*_ and *resistant*_*96h*_ groups (Tukey’s HSD; respectively *t* = −3.28; *P* = 0.01 and *t* = −3.42; *P* < 0.01). However, no difference was observed between *control*_*96h*_ and *resistant*_*96h*_ groups (Tukey’s HSD; *t* = 0.12; *P* = 0.99).

**Figure 2:**
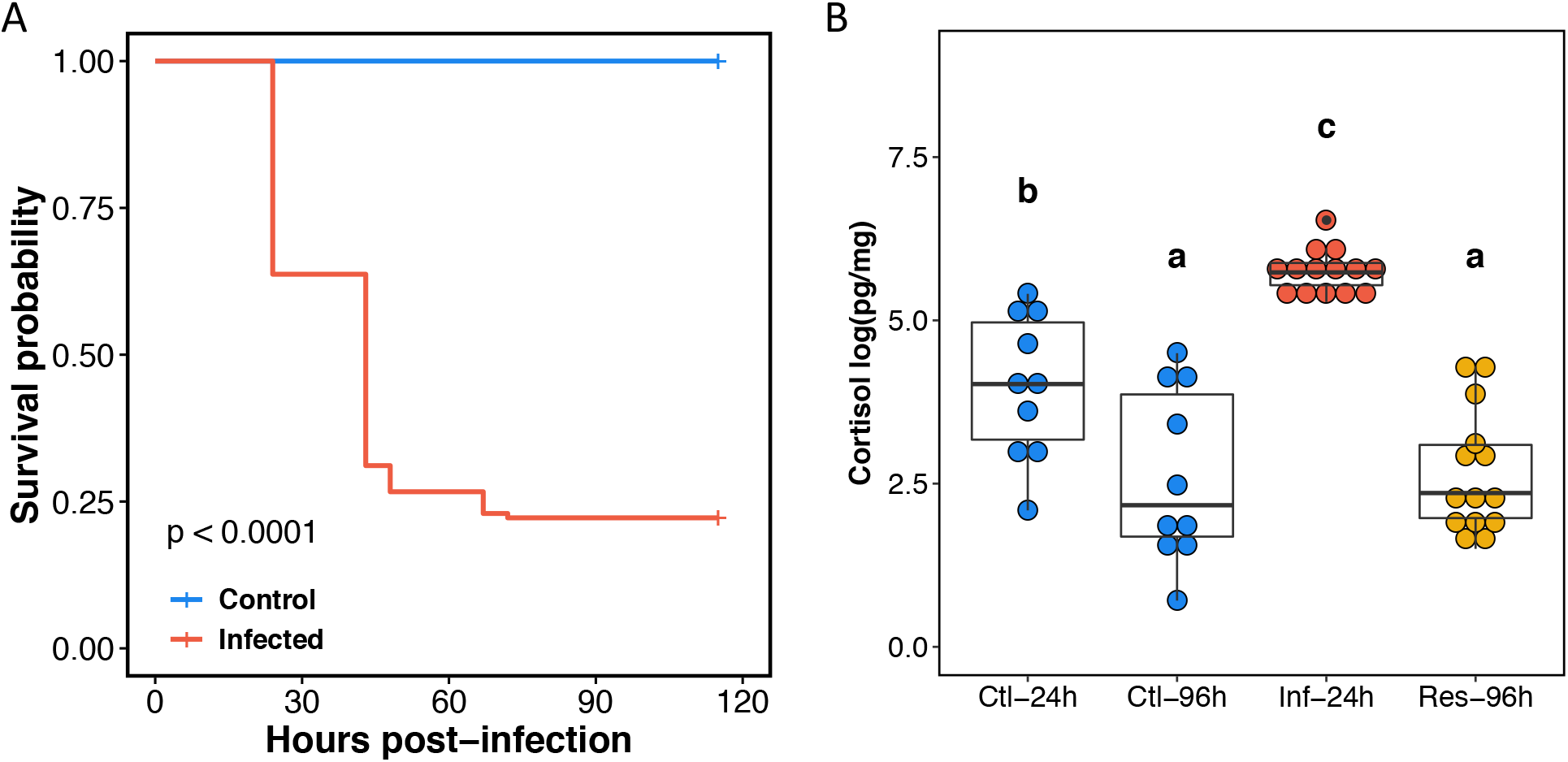
Kaplan Meier survivals estimates and scales cortisol levels. **A)** Kaplan–Meier survival curves for control (blue) and infected (red) groups over the 115hpi of the experiment. Survival was monitored at 0,19, 24, 43, 48, 67, 72, 91, 96 and 115hpi. **B)** Scale cortisol levels are expressed on a logarithmic (Log10) scale. Ctl-24h: *control*_*24h*_; Ctl-96h: *control*_*96h*_, Inf-24h: *infected*_*24h*_; Res-96h: *resistant*_*96h*_, groups. Letters represent significant differences, *P* < 0.05, Tukey’s HSD test.

### b) Dynamic of host transcriptomic response to infection and search for genomic bases of resistance

Global mean unique mapping rate for skin smear samples reached 71.64 ± 2.99% against a combined reference for host and microbes compartments. Datasets were predominantly composed of host-origin sequences (mean 86.57 ± 13.48%), with *infected*_*24h*_ group showing significantly higher proportion of non-host origin reads [mean 30.70 % ± 0.03 se] than other groups (Dunn’s test; Benjamini-Hochberg adj. *P* < 0.05; Figure S1). Details of host’ transcriptome and individuals mapping are provided in Table S1 and S3.

**Figure S1:**
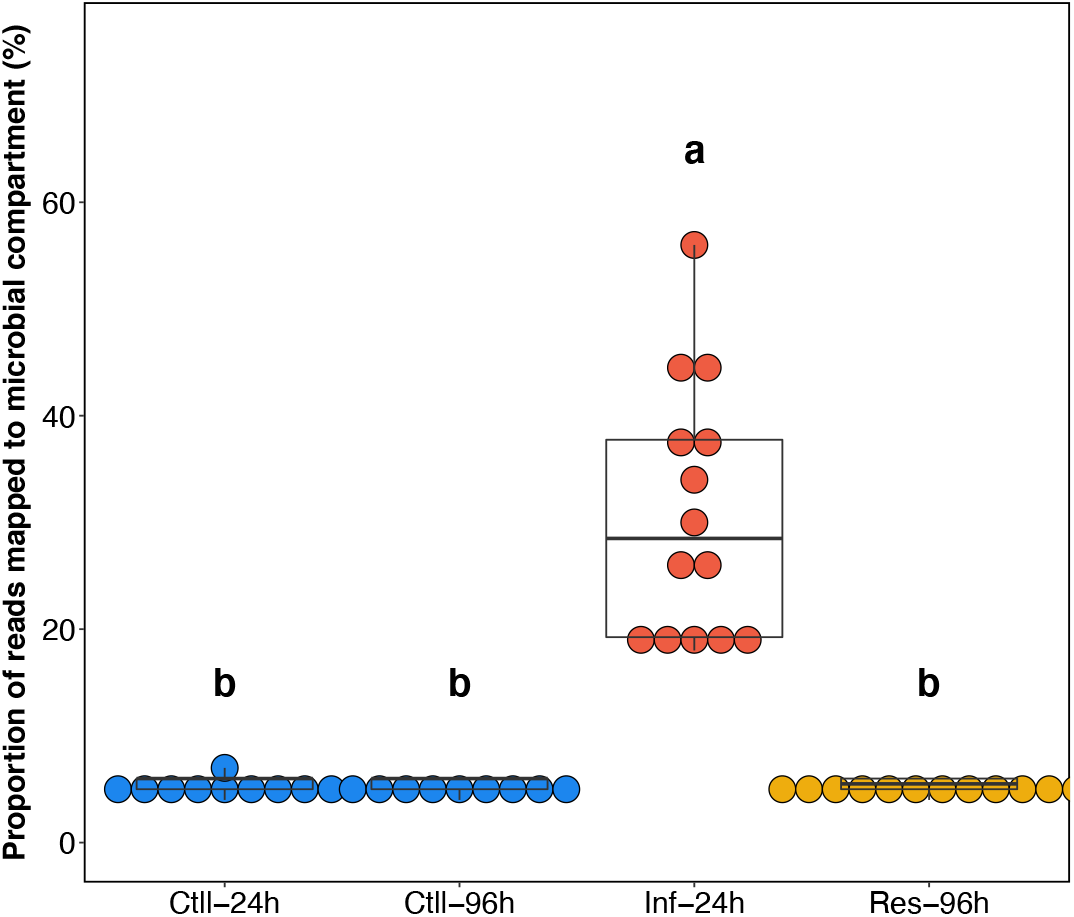
Proportion of the reads mapped to the microbial compartment. Reads origin was dissociated *in silico*. Reads were considered originating from microbial compartment when no mapping was evident in the fish transcriptome. Ctl-24h: *control*_*24h*_; Ctl-96h: *control*_*96h*_, Inf-24h: *infected*_*24h*_; Res-96h: *resistant*_*96h*_, groups. Letters represent significant differences, *P* < 0.05, Dunn’s test.

#### Fish response to infection

Differential expression analyses reveal strong differences in host genes expression profiles between *control*_*24h*_ and *infected*_*24h*_, with a total of 3,631 and 2,388 down and up-regulated genes in *infected*_*24h*_, respectively, compared to *control*_*24h*_, (|FC| > 2; FDR < 0.01; Table S4). *Infected*_*24h*_ group responds to infection mainly by activating immune system response (Biological Process; BP), sterol biosynthetic process (BP), defense response (BP), inflammatory response (BP), regulation of biological quality (BP), lipid metabolic process (BP), iron ion homeostasis (BP), complement binding (Molecular Function; MF), heme binding (MF), oxidoreductase activity (MF), sulphur compound binding (MF), (1->3)-beta-3-D-glucan binding (MF). A complete list of GO enrichment for each module is provided in Table S4.

We further used co-expression network analysis (WGCNA) to draw clusters of co-regulated genes associated with discrete (condition) or continuous variable (weight and cortisol) and to identify putative hub genes. No genes module significantly correlates with fish mass suggesting that it had no significant effect on genes expression profiles. A total of three modules show negative correlation (*P* < 0.01) with disease status (coded 1 for *control*_*24h*,_ *control*_*96h*_ and *resistant*_*96h*_ groups and 2 for i*nfected*_*24h*_) namely module_turquoise-host_ (r = −0.97, *P* < 0.001), module_black-host_ (r = −0.5, *P* < 0.001) and module_green-host_ (r = −0.49, *P* < 0.001; Figure 3). Inversely, two modules show positive correlation with the condition namely module_blue-host_ (r = 0.97, *P* < 0.001) and module_pink-host_ (r = 0.48; *P* = 0.001). Almost all these modules (with the exception of module_black-host_) also correlate significantly with cortisol levels. The genes found up-regulated in *control*_*24h*_ clustered mostly in the module_turquoise-host_ (n = 3,468; 95.5%), module_black-host_ (n = 72; 2.0%) and module_green-host_ (n = 39; 1.0%). Nearly all the genes found up-regulated in *infected*_*24h*_ clustered in module_blue-host_ (n = 2,352; 98.5%). Not surprisingly, main drivers genes (‘hub-genes’) encompass several transcriptional activators such as for the module_turquoise-host_ several Zinc finger proteins, Transcription factor GATA-3 (*gata-3*), Forkhead box protein O3 (*foxpo3*), activators of the autophagy pathways and main drivers of naïve specific T-cells differentiation and activation [56], Runt-related transcription factor 2 (*runt2*) coding genes, involved in osteoblast differentiation, a mineral depositing cells and enhancer of T-cells receptor and sialoproteins [57]. In the module_blue-host_, ‘hub-genes’ mainly report actors of the innate immune system, inflammatory response, wound healing, oxidative and adhesion activity (Figure 3B).

**Figure 3:**
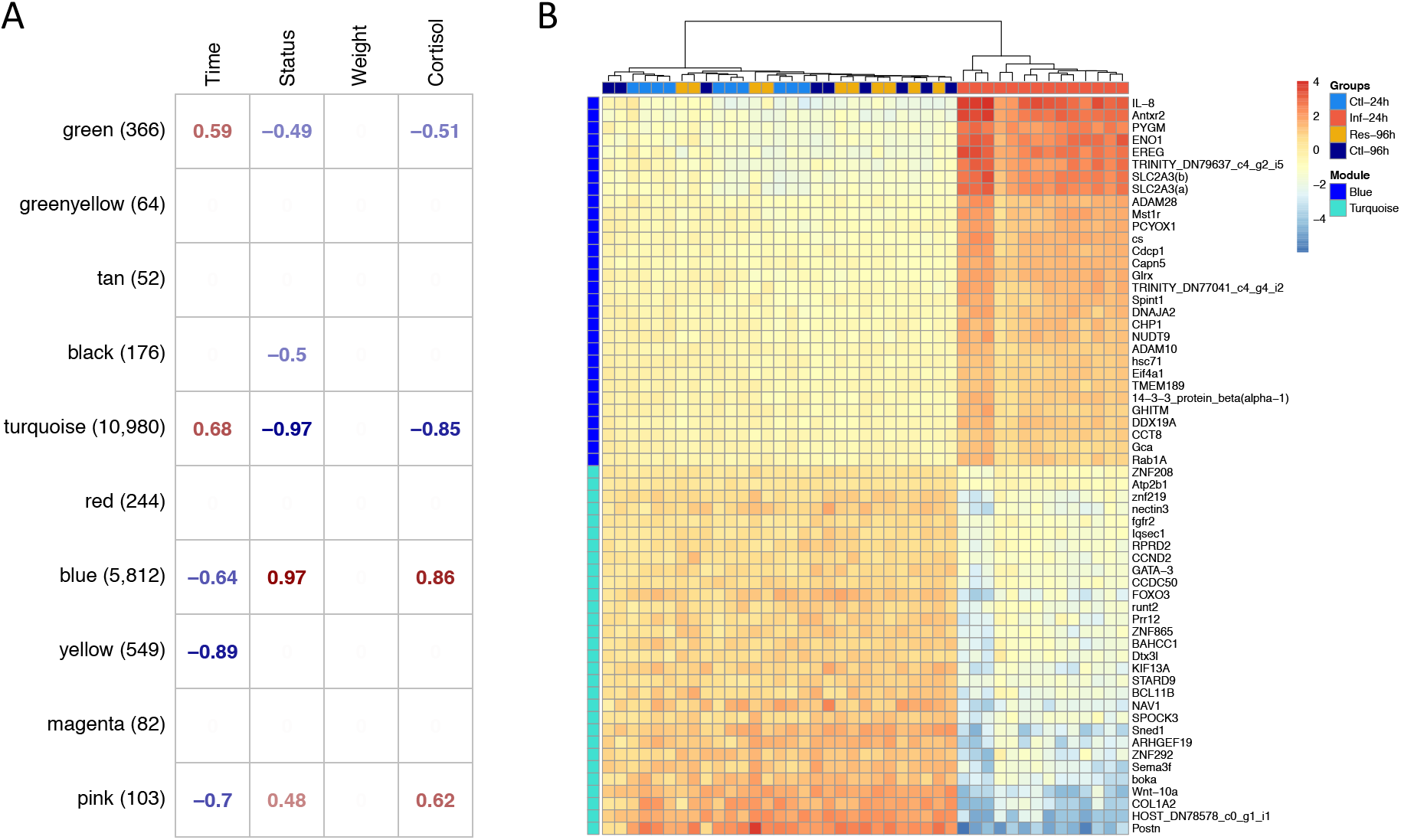
Signed co-expression network analysis for *P. orbicularis*. A) Correlation matrix for *P. orbicularis*. Values in the cells represent significant (*P* < 0.01) Pearson’s correlation of module eigenvalue to physiological parameters (top panel). Names (left panel) are arbitrary color-coded names for each module; values in parenthesis represent the number of genes per module. Empty cells indicate non-significant correlation (*P* ≥ 0.01). Individuals’ cortisol [log(pg.mg^−1^)] and weight (g) are continuous values. Time (24hpi and 96hpi) and Status (coded 1 for *control*_*24h*,_ *control*_*96h*_ and *resistant*_*96h*_ groups and 2 for i*nfected*_*24h*_) are discrete numeric values. B) Heatmaps of top 30 genes in module_blue-host_ and module_turquoise-host._ Scales represent Log2 (prior 2) of the individuals’ expression levels. Individuals were clustered using hierarchical clustering procedures implemented in pheatmap R package **[58]**.

#### Genomic bases of resistance

We found 38 DEGs between *control*_*96h*_ and *resistant*_*96h*_ (16 and 22 down and up-regulated in *resistant*_*96h*_, respectively; |FC| > 2; FDR < 0.01). GO analyses show that adaptive immune response (BP) tends to be activated (uncorrected *P* < 0.001) in *resistant*_*96h*_ that includes genes related to pathogens recognition and immune response such as C-type lectin domain family 4 member M, Low affinity immunoglobulin gamma Fc receptor II-like and T-cell receptor beta variable 7-2 coding genes. Inversely, *resistant*_*96h*_ show inactivation of the regulation of ERK1 and ERK2 cascade (BP) and repressors of the response to wounding (BP), regulation of the transforming growth factor-beta secretion (BP), alcohol biosynthetic process (BP) and regulation of interleukin-8 production (BP). However, GO enrichments were not considered significant under our threshold (Bonferroni adj. *P >* 0.05). A total of 27 (71.1%), out the 38 DEGs identified, also showed different expression levels between *control*_*24h*_ and *infected*_*24h*_. Among the 11 remaining genes (28.9%), we found Arf-GAP with dual PH domain-containing protein 1, C-type lectin domain family 4 member M, protein KIAA1324-like homolog, Ankyrin repeat and fibronectin type-III domain-containing protein 1 and T-cell receptor beta variable 7-2, up-regulated in *resistant*_*96h*_ group. Inversely, we found the Sal-like protein 1, Early growth response protein 1 and the Low affinity immunoglobulin gamma Fc receptor II-like down-regulated in *resistant*_*96h*_. The complete list of DEGs and GO term enriched is provided in Table S4.

We finally searched for genetic variation (SNPs) across *resistant*_*96h*_ and *infected*_*24h*_ (two established phenotypes) in order to identify putative variants associated with resistance capacities. We identified a subset of 13,448 filtered bi-allelic SNPs. Genetic variations analyses did not suggest any significant difference among groups (relatedness, Fst) and was not correlated with any of the groups’ cortisol levels or fish mass based on the 13,448 markers (PERMANOVA; 1000 permutations; *P* = 0.18; Figure S2).

**Figure S2:**
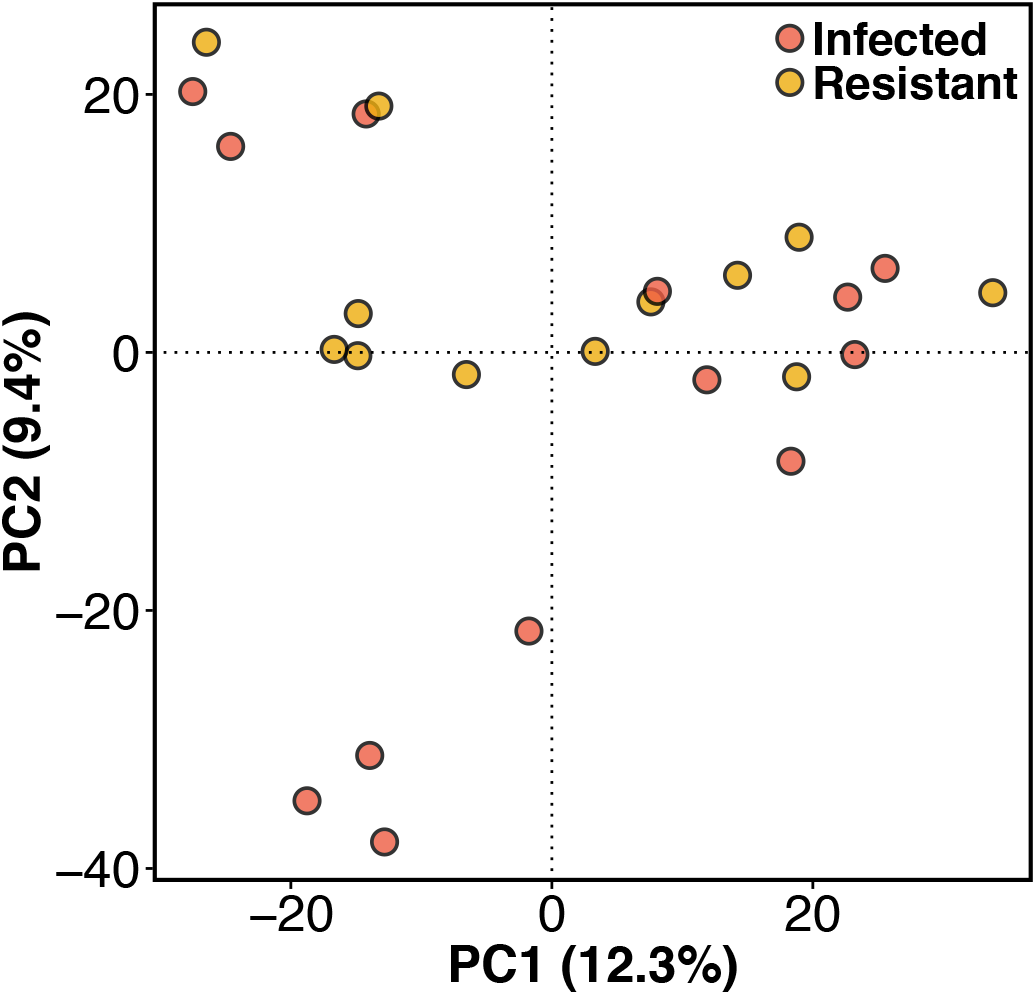
Genetic variation and relatedness across established phenotypes. Principal component analysis for the total filtered dataset including 13,448 bi-allelic markers.

### c) Microbiome flexibility and interaction among pathogens species and host response

#### Dynamic of microbiota communities on fish skin

MiSeq sequencing strategy with amplification of the 16S rRNA V4 region of the 16S rRNA resulted a mean number of PE of 231,164.02 ± 36,542.99 sd out of which a mean of 82.47 ± 2.54 remained after filtering (Table S2). Species richness (Shannon) was lower in *infected*_*24h*_ than other groups (MWW; Holm adj. *P* < 0.001). Similarly, *control*_*24h*_ shows reduced species diversity values compared to *control*_*96h*_ (MWW; Holm adj. *P* < 0.001) but no difference was observed between *resistant*_*96h*_ and *control*_*96h*_ (MWW; Holm adj. *P* = 0.71; Figure S3A). Similar observation was made for Fisher’s alpha parameter but the latter shows significant difference between *control*_*24h*_ and *resistant*_*96h*_ (MWW; Holm adj. *P* = 0.05; Figure S3B). Bacterial communities vary significantly across groups (PERMANOVA; F = 30.93; *P* < 0.001); however, the beta dispersion also differs significantly (‘betadisper’, ANOVA, F = 30.93; *P* < 0.001). Specifically, based on Bray-Curtis distances values, communities are less variable in *infected*_*24h*_ group compared to all other groups (Tukey’s HSD, [*control*_*24h-*_*infected*_*24h*_: 95% CI= 0.05-0.19; *resistant*_*96h*_*-infected*_*24h*_: 95% CI= 0.17-0.31; *control*_*96h*_ *-infected*_*24h*_: 95% CI= 0.12-0.28]; *P* < 0.001, Figure S4). Communities are also more variable in *resistant*_*96h*_ compared to *control*_*24h*_ (Tukey’s HSD, 95% CI= 0.04-0.20; *P* = 0.001). The ASVs associated with *Tenacibaculum (Flavobacteriales)* are largely enriched in *infected*_*24h*_ compared to *control*_*24h*_, but also significantly enriched in *resistant*_*96h*_ compared to *control*_*96h*_ (shrinked |Log2FC| > 2; FDR < 0.01; Figure S4).

**Figure S3:**
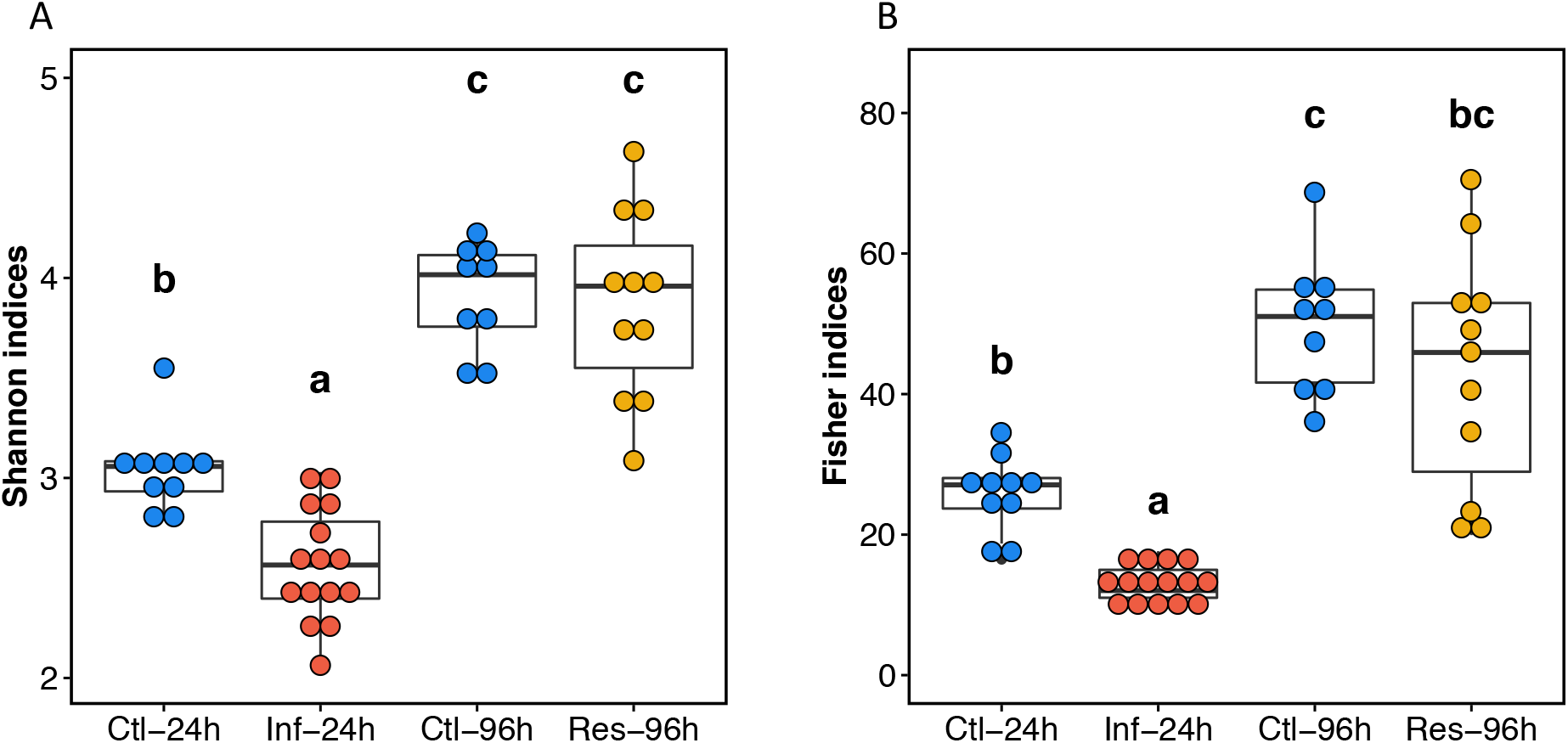
Alpha-diversity estimates across groups. Alpha-diversity was computed using A) Shannon (H’) and B) Fisher indexes. Ctl-24h: *control*_*24h*_; Ctl-96h: *control*_*96h*_, Inf-24h: *infected*_*24h*_; Res-96h: *Resistant*_*96h*_, groups. Letters represent significant differences, *P* < 0.05, Tukey’s HSD. Each dot represents an individual.

**Figure S4:**
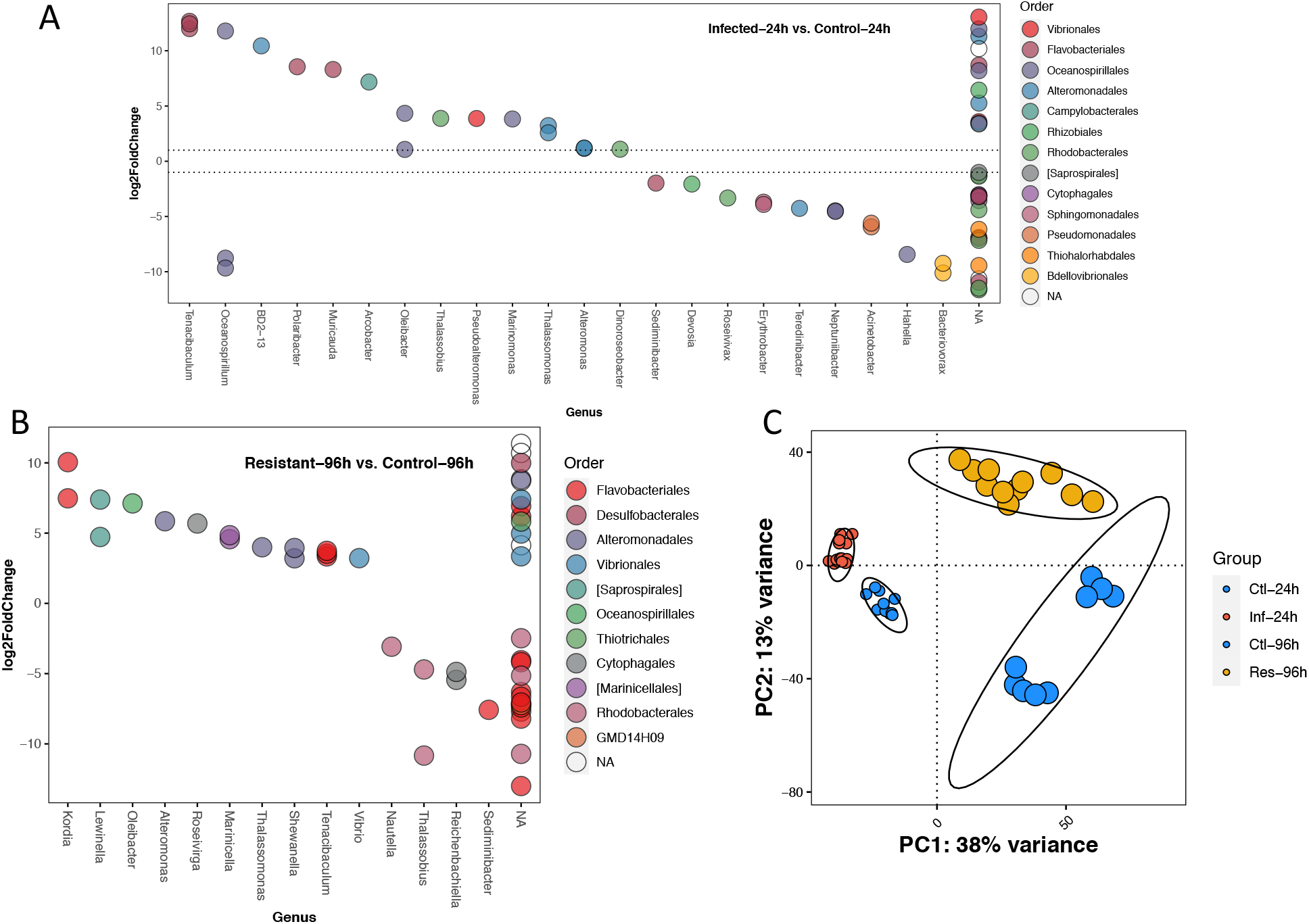
Taxa enrichment and beta-diversity dissimilarities across groups. A) ASVs enrichment between *infected*_*24h*_ (positive log2FC) and *control*_*24h*_ (negative values). Colors represent different Orders. The y-axis reports shrinked Log2FC. Horizontal dash lines represent Log2FC threshold for significance (|Log2FC| > 2). NAs represent missing taxonomic information for this ASV; B) ASVs enrichment between *resistant*_*96h*_ (positive log2FC) and *control*_*96h*_ (negative values). Colors represent different orders. The x-axis represents associated Genus, y-axis reports shrinked Log2FC. NAs represent missing taxonomic information for this ASV. C) PCoA based on Bray-Curtis distance values. Ellipses represent 95% CI intervals. Ctl-24h: *control*_*24h*_; Ctl-96h: *control*_*96h*_, Inf-24h: *infected*_*24h*_; Res-96h: *Resistant*_*96h*_, groups. Smaller points code for 24hpi, larger points for 96hpi groups.

#### Long-reads refinement of bacteria communities in infected fish

Results from Nanopore sequencing on full 16S rRNA sequences served to refine the taxonomy at the species levels, which might be limited with short-reads approaches. We amplified the full 16S rRNA for 8 individuals randomly subsampled from the infected group 24hpi resulting in a mean number of SE reads of 60,019.62 ± 33,778.99 sd after pre-processing (individual with the lowest coverage reached a total of 29,520 sequences). Nanopore shows an over-dominance of *T. maritimum* in *infected*_*24*_ (73.51 ± 4.89%) and confirmed the the presence of other genera, including *Vibrio, Polibacter, Alteromonas* and *Pseudoalteromonas* (Figure S5). Among these genera, some species are known as potential fish pathogens such as *V. harveyi* [59], while others (*Pseudoalteromonas*) have been proposed as putative probiotics [60].

**Figure S5:**
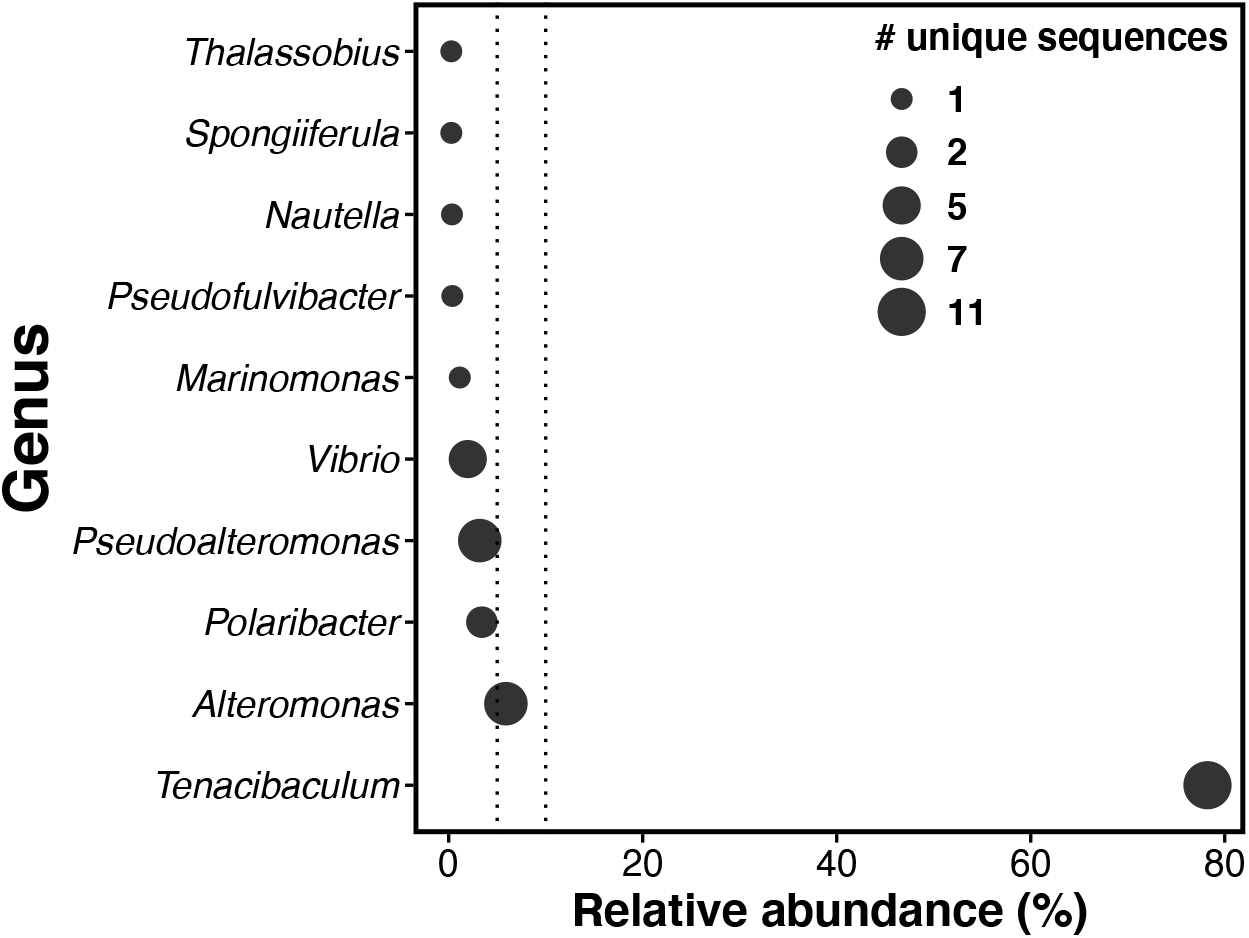
Relative abundance of bacterial genera in the *infected*_*24h*_ group reported with Nanopore. 16S rRNA sequences and their relative abundance were generated from Nanopore reads (see methods). These sequences were blast against NCBI nt database (BLASTn; e-value<10-5). Size of the dots represent the number of unique sequence per genus.

### d) Microbial compartment transcriptomic activity

#### Genes expression of *Tenacibaculum maritimum* during experimental infection vs *in vitro*

We explored the genes expression levels for *T. maritimum* in the fish during the pic of infection compared to *in vitro* to highlight putative genes associated with virulence (Figure 4). Mean total mapped reads against *T. maritimum in vivo* reached 5.65M ± 0.82 se (Figure S6), which is sufficient to conduct differential expression analysis [61].

**Figure 4:**
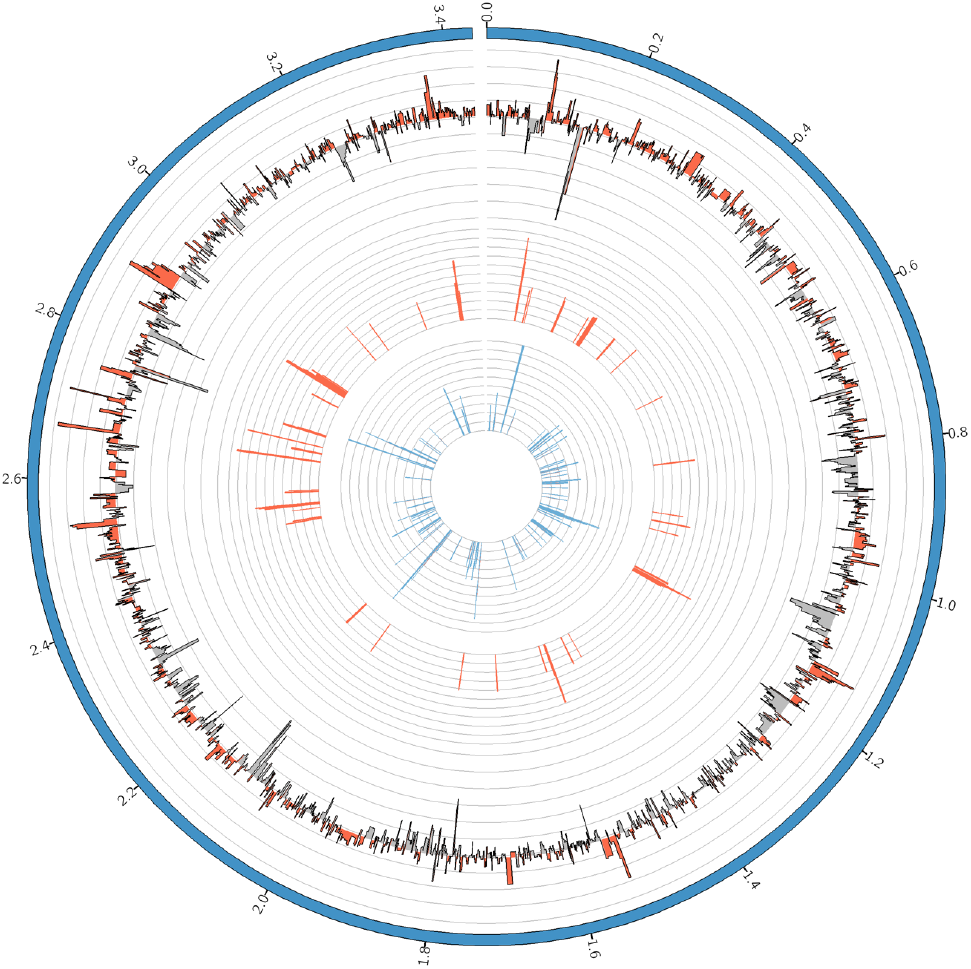
Circos plot of *in vitro* and *in vivo T. maritimum* expression comparisons. External line represents mean shrinked log2FC *in vitro* (negative values) compared to *in vivo* (positive values). Reds bar represent shrinked log2FC of significantly up-regulated genes *in vivo*; blue histograms represent shrinked log2FC of significantly up-regulated genes *in vitro*. Circos positions were based on *T. maritimum* NCIMB 2154^T^ genome information. Positions represented by external ticks are reported in million bp.

**Figure S6:**
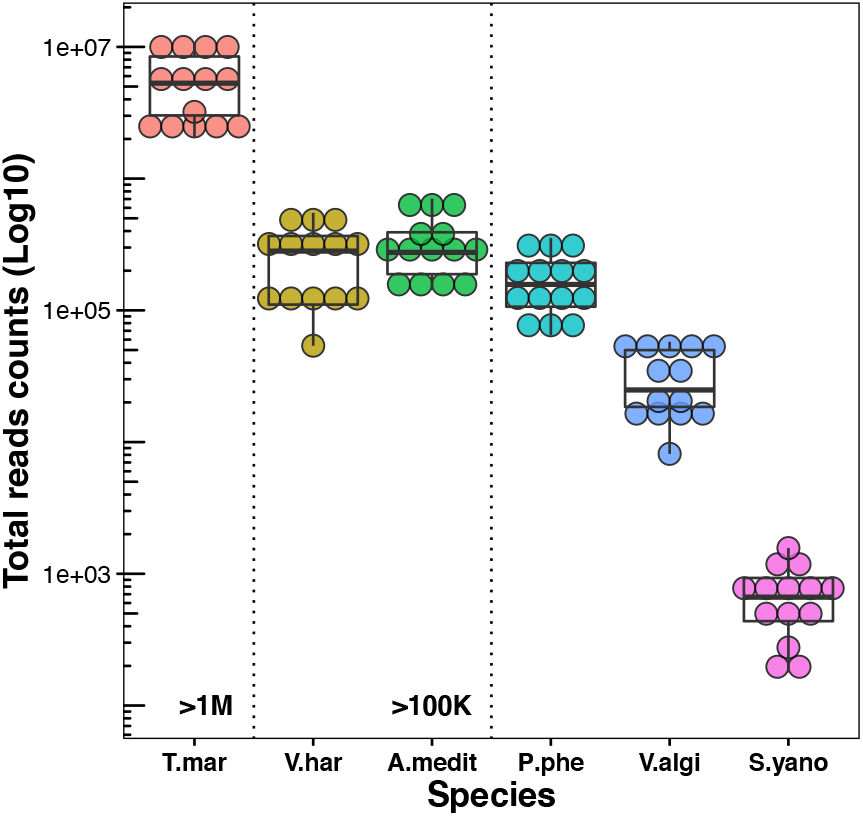
Total Illumina PE reads count for the most abundant species in the microbial compartment. The selected reference species were the most abundant species represented in the Nanopore 16S rRNA analysis (see methods). T.mar: *T. maritimum*; V.har: *Vibrio harveyi*; A.medit: *Altermonas mediteranea*; P.phe: *Pseudoalteromonas phenolytica*; V.algi: *Vibrio alginolyticus*; and S.yano: *Sphingobium yanoikuyae*. Each dot per species represents one individual.

We found a total of 72 and 142 DEGs up-regulated during experimental infection (*in vivo*) and *in vitro*, respectively (Shrinked |log2FC| > 2; FDR < 0.01). We only found the sulfate assimilation (BP) function enriched *in vitro* (Bonferroni adj. *P* < 0.1). Among the GO positively enriched during experimental infection (Bonferroni adj. *P* < 0.1) we found the glucan catabolic process (BP), external encapsulating structure part (cellular component, CC), pattern binding (MF) and the antibiotic catabolic process (BP). These processes include genes already highlighted in genome scale comparisons of *Tenacibaculum* species as possible virulence-related factors such as the catalase/peroxidase *katG* or several Ton-B receptor dependent and Suc-C and SucD-like proteins [19] (Figure S7). In parallel, we also detected putative candidate involved in host membrane interaction and integrity such as Ulilysin, Streptopain, Pneumolysin toxin and peptidoglycan-associated factor lipoprotein or adhesines. A complete list of GO is provided in Table S4.

**Figure S7:**
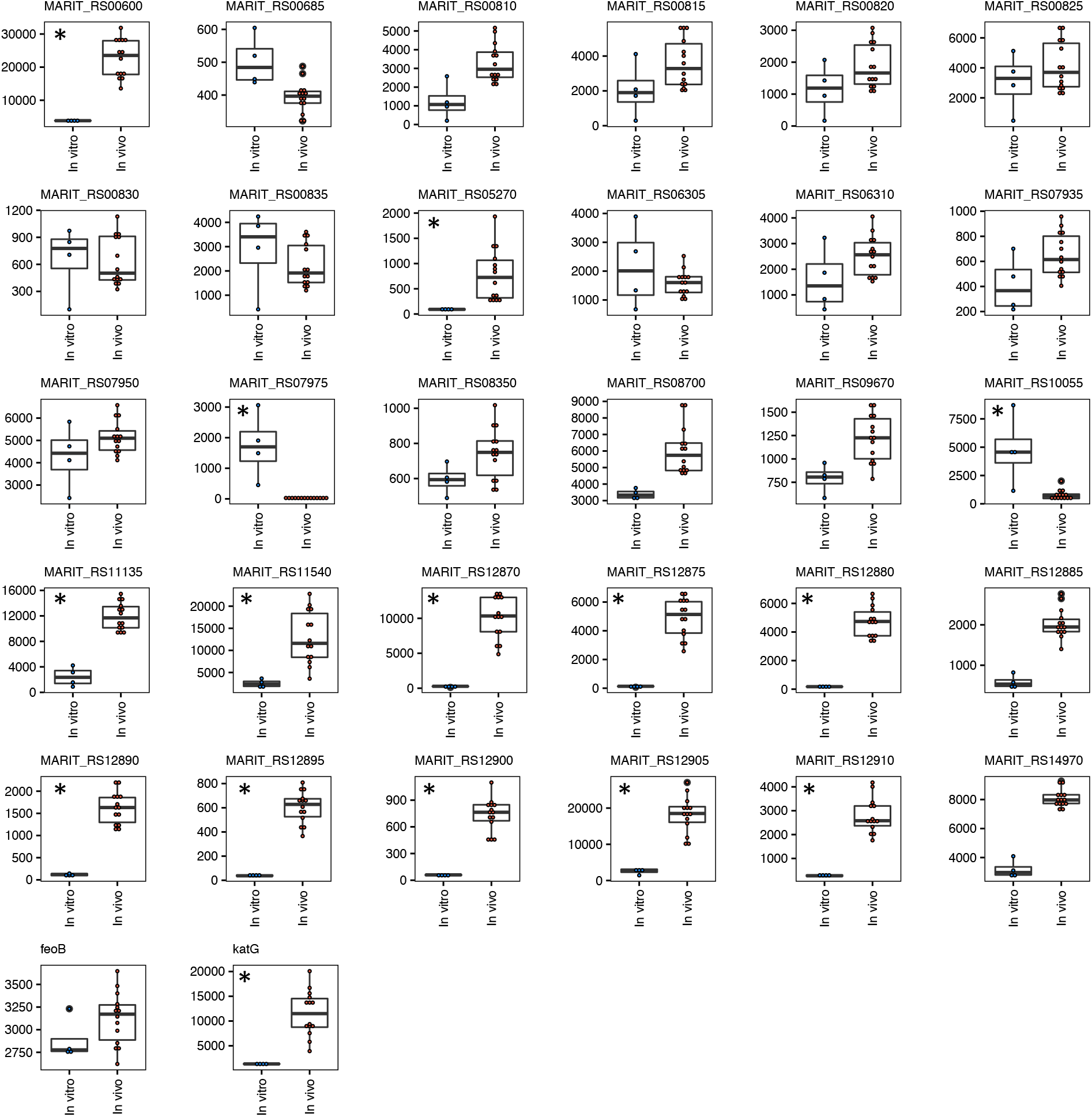
Plot of expression levels between *in vitro* and *in vivo* groups for all the virulence-related genes previously identified. The gene list was obtained from whole-genome analysis in *Tenacibaculum* spp. [19]. Asterisks represent genes with significant difference between groups (Shrinked |Log2FC| > 2; FDR < 0.01).

## Discussion

Tenacibaculosis is a worldwide fish disease responsible for considerable farmed fish mortality events; yet, knowledge is still missing on the microbiome kinetics during infection and the concomitant host-pathogens interaction. Dual RNAseq arose as a method of choice by providing unprecedented simultaneous information on the molecular features of the infection. It is particularly adapted for systems characterized by a massive pathogens burden with readily accessible material and for which cultures are not available [17,62]. Here we adapted this approach for tenacibaculosis in *P. orbicularis* fish skin samples with the goal to comprehensively assess genomic basis and kinetics of the infection as well as associated resistance mechanisms.

### Infection modulates a wide range of host’ innate and adaptive immune effectors

Bath exposition of *T. maritimum* was highly efficient in inducing tenacibaculosis in juvenile orbicular batfish. The low survival rate and the kinetics of infection support what is usually observed for other fish species [16,20,63]. Infected fish, sampled at the pic of infection (24hpi), show large skin lesions characteristic of tenacibaculosis together with high cortisol concentration in their scales [32]. Cortisol mediates changes in individual energy balance (e.g. mobilisation of energy stores; immunity; cognition; visual acuity; and behaviour) [64,65]. This initial cascade of physiological and behavioural changes enable the organism to cope with acute stressors by mobilising adequate bodily functions, while concurrently inhibiting non-essential functions (e.g. reproduction, digestion) [66]. Here increasing cortisol levels reflect local stress response to unfavourable environment and is most likely involved in triggering fish rapid immune response [67].

As expected fish immune response, especially innate immune system is strongly solicited at 24hpi. Infected fish show activation of the acute inflammatory response, mainly through drivers genes including interleukin-8 (IL-8) [68], but also activation of pathogens recognition receptors (PRRs), chemokines as well as antimicrobial related humoral effectors. For instance, infection triggers co-expression of the cascade Toll-like receptor 5 (TLR5) and Myeloid differentiation primary response protein (MyD88), as previously reported in bony fish during bacterial infection [69]. However, the diversity of the fish immune actors along with a relatively limited knowledge on specific effectors functions significantly hampered the comprehensive understanding of the mechanisms involved in our non-model species. For instance, in parallel of the TLR5, several other TLRs show reduced expression in infected fish, including TLR2 type-1, TLR-8 and non-mammalian (‘fish-specific’) TLR21. Despite previous effort toward assessing diversity of TLRs sequences, protein-specific function remains poorly known in teleost [70]. Similar observations can be made for the Complement system, specifically complement C3, a key component of the immune system involves in “complementing” antibodies for bacteria cells killing [71], for which several isoforms are reported in the platax transcriptome. The different isoforms here have divergent patterns of expression (both up or down-regulated in *infected*_*24h*_ group), which support previously observed difference in target surface binding specificities [72].

Innate immune response is generally tightly linked with cellular homeostasis regulation and precedes adaptive immune response. The ability of the fish to maintain cellular homeostasis during infection is of primary importance when facing infection and mechanisms include redox, biological quality control (autophagy) as well as ion levels maintenance [73,74]; all found affected in the platax. For instance, infected fish largely activates effectors of the iron ion homeostasis. Iron, albeit largely present in the environment, is poorly accessible by organisms and iron sequestration and maintenance is a major mechanisms developed by the host to limit pathogens growth as well as to regulate macrophages cytokines production [75]. In parallel, *infected*_*24*_ activate the (1->3)-beta-D-glucan binding process and contribute to the body of literature revealing receptor capacities in fish and pathways conservation (through PRRs, C-lectin and/or TLRs) across vertebrates and invertebrates [76,77]. Indeed, supplementation of β-glucan stimulates immune response in fish and confers higher resistance of the host to virus and pathogens (probably by reducing bacteria adhesion through lectin binding [78]); hence representing promising immunostimulant in aquaculture [77,79]. Effect of β-glucan vary depending on the species, exposure time, source of glucan, organs and markers monitored [76,80] and further studies will be needed to evaluate its potential at the production scale. Nevertheless, β-glucan is also relevant in bridging inflammatory response and activation / differentiation of T-cells in the adaptive immune response [81].

The adaptive immune response was also modulated at 24hpi and its fine-tuned orchestration offers the opportunity to dissect preferential immune paths to fight against *T. maritimum* infection. We identified several hallmarks of differentiated T-cells, indicative of the specialisation of the adaptive immune response to *T. maritimum* infection. Among the main driver genes of the response to infection in platax we noted a reduced expression of *foxp3* and *gata-3* in *infected*_*24h*_. Both transcription factors are important regulators of Naïve CD4+ naïve T-cells fate encouraging differentiation to T-regulatory (Treg) [82] and T-helper 2 (Th2) cells [56], respectively. Similarly, we show reduced expression of T-bet transcription factors, an hallmark of Th1 cells [56]. Inversely, infected fish seem to activate Th17 cells differentiation as suggested by simultaneous activation of signal transducer and activator of transcription (STAT1-alpha/beta) and cytokine IL-17 [82]. The Th17 cells are mainly dedicated to control bacterial and fungi entry [83]. Our results suggest and support previous a complex orchestration of T-cells differentiation via antigens communication and associated cytokines regulatory network in platax during *T. maritimum* infection [56,84]. However, we can’t over rule that changes in transcripts abundance might also be indicative of cells migration and certainly, other approaches including cellular imaging [85] would clarify the presence and refine the regulation of T-reg cells in fish.

### The genomic bases of resistance in *P. orbicularis*

Despite profound activation of the immune system response, most of the fish failed at containing the infection. At 96hpi, less than 25% of the infected fish had survived the bacterial challenge. These surviving individuals did not display any skin lesion, suggesting they resisted the *T. maritimum* penetration and/or limited initial bacterial adhesion. Considering the high bacterial concentration in the tanks during infection and the severity of the mortality event, it is very unlikely that resistant fish might have totally escaped contact with the pathogen. Indeed, *T. maritimum* were present in *resistant*_*96h*_ but not in *control*_*96h*_ (see next section for details). Fish were thus able to maintain the integrity of first barrier against pathogens; hence we hypothesize that genes activity difference would reveal specific immune actors present and/or efficient enough (timing of regulation) to inhibit pathogens multiplication and entry in resistant fish [16]. We show that PRRs, specifically a C-type lectin receptor, was up-regulated in *resistant*_*96h*_, together with a T-cell receptor and Low affinity immunoglobulin gamma Fc receptor II-like while fibronectin coding genes was down-regulated. These genes-products are known for binding, agglutinating and neutralizing bacteria [86] as well as triggering humoral immune response [87] or providing extracellular structure for pathogens adhesion through fibronectin-binding proteins [88,89]. Our experimental design did not allow segregating between a basal difference in expression in *resistant*_*96h*_ (genomic basis of resistance *per se*) or if the difference at 96hpi is the result of a delayed adaptive immune response (timing of genes expression). Nonetheless, in catfish, lectin expression allows differentiating resistant from susceptible families against *Flavobacterium colummnare*, another gram-negative bacteria of the *Flavobacteriaceae* family [90]. Further longitudinal studies monitoring genes expression of resistant fish along the entire infection (and before the challenge) should prove useful in identifying resistant-specific response to infection. Ideally, these studies should also simultaneously look at different immune-specific organs and tissues compartments and integrate genome-scale genetic variants (not limited to coding regions) to infer putative genetic bases of resistance.

### Microbiome dynamics and host-pathogens communication

At 24hpi, microbiome community was dramatically affected with over-dominance of *T. maritimum*. Abundance of *T. maritimum* was evident from metabarcoding data and contributes to significantly reduce species richness in *infected*_*24h*_ fish. We went further and compared expression of *T. maritimum in vivo* (during infection) compared to *in vitro*, with the hypothesis that key drivers of pathogenicity would be solicited for thriving and breaking host’ defense barriers.

There are at least two major challenges that *T. maritimum* needs to overcome to successfully infect the host: Pathogens need 1) to compete (at the intra and interspecific levels) for resources to metabolise from the local environment and 2) to resist to the host immune responses and stressful conditions. During infection *T. maritimum* enhances its glucan catabolic activity; while this might only reflect differences due to changes of the local environment conditions (host mucus and skin) and/or resources availability, it also reveals some major mechanisms explaining the success of *T. maritimum* to grow on fish skin. Among the genes involved in glucan catabolic, we report several key components linking alternative food and minerals supplies and putative virulence-related functions, such as several genes involve in specifically degrading and up taking sialoglycan and ions, mainly iron [19]. Sialidase activity explains *Capnocytophaga canimorsus* burst when in contact with host cells it allows the pathogen to mobilise sugar directly from host phagocytes [91]. Similarly, the tonB coding gene, regularly reported as a gene relevant for pathogenicity, confers virulence to *Edwardsiella ictaluri* by allowing to maintain growth in iron-depleted medium [92]. In parallel, several stress resistance related genes were activated during experimental infection; all also involved in the antibiotic catalytic functions. These genes include *katA* and *katG*, coding for two catalases-peroxidases involved in resistance to reactive oxygen species (ROS), by detoxifying exogenous H_2_O_2_ produced by host macrophages as a defense mechanism [93]. Obviously, the identification of virulence-related genes can not be limited to the ones differentiating *in vitro* versus in vivo infectious status and other actors might be involved in conferring *T. maritimum* its pathogenicity. For instance, siderophore coding genes are constitutively expressed *in vitro* or during experimental infection. These genes are a determining factor of host-pathogens and pathogens-pathogens interactions in the so-called “race for iron” [94–96], which suggest that *T. maritimum* is highly efficient in mobilizing iron, independently of the local environment.

Finally, we mostly explored here expression levels variation in the light of an exclusive interplay between host and *T. maritimum*, which might be effective considering the over-dominance of *T. maritimum* in fish mucus. However, most of the infection systems report several pathogen co-occurrences and the presence and/or activity of other opportunistic pathogens might also play an important role in host’ fate [15]. In infected fish, we found a relatively large representation of those so-called “opportunistic” bacteria, known for their pathogenicity in fish, including *V. harveyi. V. harveyi* is an ubiquitous bacteria and one of the most common pathogens inducing major disease outbreaks in fish farming [59]. The enrichment for Vibrio ASVs in *resistant*_*96h*_ mucosal communities and the absence of evident physiological associated changes (cortisol, mortality, skin integrity) in this group, suggest that *V. harveyi* alone is not sufficient to induce mortality in platax under our specific experimental conditions and associated bacterial burden.

## Conclusions

Here we provide a comprehensive description of the host and *T. maritimum* interplay under experimental infection conditions. Our results serve in deciphering the complex orchestration of innate and immune response of the host but also propose some promising avenues of research to limit the impact of tenacibaculosis in fish farming. By deploying an integrative ‘omic’ approach, we identified bacterial interaction as well as putative virulence-related genes in *T. maritimum* and candidate-genes involved in fish resistance. Importantly however, the detection of immune actors in fish and our comprehension of their regulation rely mainly on the quality of the annotation and the knowledge we have of their activity in other species, mainly mammals model species. Consequently, further studies are now urgently needed to properly apprehend genetic and genomic bases of response to infection and possible resistance capacities in other non-model fish species.

## Ethics approval

*In vivo* experiments compiles with all the sections of deliberation n° 2001-16 APF from the Assembly of French Polynesia regarding domestic or wild animals welfare, issued in ‘Journal Officiel de Polynésie française’ on February 1^st^, 2001. In the absence of *adhoc* ethical Committees, we follow European Commission, DGXI [97] and ARRIVE [98] guidelines.

## Consent for publication

Not applicable

## Data accessibility

Raw sequences are deposited in NCBI database under accession number (PRJNA656561). Codes are made publicly available on a Github repository: https://github.com/jleluyer/metatranscriptomics_workflow

## Authors’ contributions

DS and JLL conceived the experiment. MC conducted fish husbandry and rearing. DS, MC, JLL and QS conducted the sampling. CB and CB conducted DNA and RNA extractions. QS conducted the cortisol levels analyses. PA assembled the *P. orbicularis* transcriptome. JLL conducted the RNAseq and MiSeq analyses. JP conducted the Nanopore sequencing. JLL and QC conducted the Nanopore analyses. All co-authors made substantial revisions to the manuscript.

## Competing interest

The authors declare no competing interest

## Funding

The study was conducted under the Capamax project (Politique de site Ifremer) attributed to DS and JLL with financial support from Aqua-Sana convention [Ifremer-Direction des Ressources Marines (DRM)] for fish rearing and maintenance attributed to DS.

## Acknowledgement

We thank the Direction des Ressources Marines and Chedia Hamouna for their help with fish rearing. We thank Céline Reisser for providing additional gonads tissues for the platax transcriptome assembly. We also thank Eric Duchaud and Pierre Boudinot for providing valuable comments on the manuscript.

## Notes

### Competing Interest Statement

The authors have declared no competing interest.

