## Supplemental material for "Dual RNAseq highlights the kinetics of skin microbiome and fish host responsiveness to bacterial infection"

[2.] MARBEC, Univ. Montpellier, Ifremer, IRD, CNRS, F-34200, Sète, France

[3.] Virologie et Immunologie Moléculaires, Institut National de la Recherche Agronomique, Université Paris-Saclay, Jouy-en-Josas, France

[4.] Univ Polynésie française, Ifremer, IRD, Institut Louis-Malardé, EIO, F-98702 Fa'a, Tahiti, Polynésie française, France

[5.] Génomique Métabolique, Genoscope, Institut François Jacob, CEA, CNRS, Univ Evry, Université Paris-Saclay, 91057 Evry, France

[6.] USR 3278 CRIOBE, EPHE-UPVD-CNRS, Univ. de Perpignan, France

**Keywords:** Microbiome – Gene expression – 16S rRNA – Nanopore – *Tenacibaculum maritimum* – Co-infection

### Supplementary Methods & Results

#### a) Meta-transcriptomics and metabarcoding sequencing

**Dual RNA-sequencing.** Mean number of PE raw reads reached ± out of which a mean of 82.47 ± 2.54 remained after filtering (Table S1). After removing reads mapping to non-bacteria domain, number of sequences per individuals reached a mean of 105,642.4 ± 2,186.2 sd.

**Short-reads 16S rRNA metabarcoding sequencing.** We amplified the V4 region for 10-15 replicate individuals per condition. Mean number of PE raw reads reached 231,164.02 ± 36,542.99 sd out of which a mean of 82.47 ± 2.54 remained after filtering (Table S2). After removing reads mapping to non-bacteria domain, number of sequences per individuals reached a mean of 105,642.4 ± 2,186.2 sd. We identified a total of 2577 ASV across the 55 individuals.

**Nanopore full 16S rRNA metabarcoding sequencing.** We amplified the full 16S rRNA for 8 individuals randomly subsampled from the infected group 24hpi in order to characterize more precisely and accurately the composition of main skin microbes. Mean number of SE reads reached 60,019.62 ± 33,778.99 sd after pre-processing (with a minimum of 29,520 sequences).

***P. orbicularis* transcriptome reference**

**Sampling, RNA extraction and sequencing.** One individual (50 dph) of *Platax orbicularis* was first sampled for pronephros (n=1), tegument (n=1), liver (n=1) and intestine tissue (n=1) following sampling of two other individuals samples for gonads (one male and one female), to build the host transcriptome. Total RNA was extracted from the TRIZOL mix, quantity/integrity and purity were validated by both Nanodrop readings (NadoDrop

Technologies Inc.) and BioAnalyzer 2100 (Agilent Technologies). RNA was dried in RNA-stable solution (ThermoFisher Scientific) following manufacturer's recommendations and shipped at room temperature to McGill sequencing platform services (Montreal, Canada). TrueSeq v2 kit (Illumina, San 260 Diego, Ca, USA) was used to prepare mRNA depleted libraries that were multiplexed (13-14 samples by lane) and sequenced on HiSeq4000 100 bp PE sequencing device. Individuals that served for the transcriptome assembly were not included in this experiment.

**Transcriptome assembly**. A total of 382.34 M PE 100 bp raw reads (mean 63.72 M ± 7.44 sd) were filtered using Trimmomatic v0.36 [1], with minimum length (36 bp), trailing and leading thresholds of 26 and 26; respectively), implemented in Trinity v2.5.1 [2]. Reads quality was assessed with FastQC v0.11.5 (https://www.bioinformatics.babraham.ac.uk/projects/fastqc/). Reads were assembled into transcripts using Trinity v2.5.1 [2] and default parameters. The raw transcriptome was then processed in order to reduce redundancy. First, open-reading frames (ORFs) for each transcript were predicted using '*LongOrfs*' function implemented in Transdecoder v5.3.0 [2, 3]. Only the transcripts containing an ORF of at least 100 amino acids were conserved. Then, only the most expressed isoform for each gene with a minimum mapping rate of 0.5 transcript per million (TPM) was conserved. Illumina adapters were screened in the transcriptome using a blastn (version 2.6.0) approach and adapter list (http://omicsoft.com/downloads/ngs/contamination_list/v1.txt). Reads were then mapped back on the filtered transcriptome to evaluate individual mapping rate with BWA mem v0.7.15. For quality checks, the *de novo* transcriptome completeness was assessed with the BUSCO v3.0.2 [4] metazoan single-copy (n=978) database. We added a final step to look for bacteria contamination by using blastn against the NCBI nt database (release 2018-08-27). Transcripts having a hit on Bacteria (e-value<$10e^{-4}$) were discarded. The resulting transcriptome was then annotated using Trinotate pipeline v3.1.1 (https://github.com/Trinotate/Trinotate.github.io) following standard guidelines. Detailed procedures for host transcriptome assembly are available in a Github repository (https://github.com/paulineauffret/Transcriptome_platax). Transcriptome statistics are provided in Table S3.

#### b) Meta-transcriptomics functional analysis

**Differential expression and Gene Ontology enrichment.** We used a combination of differential expression and network analyses to explore host and pathogens changes in gene expression profiles during and post-infection. The genes were then used for comparing functional differences based on GO enrichment analyses. Details of the results are provided in Table S4**.**

**Table S 1: Individual mapping statistics against the combination of *P. orbicularis* transcriptome and bacterial genomes.** Transcriptome column refers to samples included in the meta-transcriptome assembly. Hpi = Hours post-infection. Mapping rate values are computed based on mapping against the combined reference transcriptome (host + microbiome).

| Alias | Hpi | Tissu | Condition | PE raw reads (M) | PE filtered reads (M) | Mapping rate (%) | Relative total bact. abundance |
|---|---|---|---|---|---|---|---|
| 1 | 24 | Skin | Infected | 84.30 | 71.50 | 76.35 | 0.38 |
| 2 | 24 | Skin | Infected | 68.65 | 58.00 | 74.47 | 0.44 |
| 3 | 24 | Skin | Infected | 56.10 | 47.65 | 72.76 | 0.19 |
| 5 | 24 | Skin | Infected | 56.01 | 47.94 | 73.60 | 0.34 |
| 6 | 24 | Skin | Infected | 71.48 | 61.90 | 73.28 | 0.30 |
| 7 | 24 | Skin | Infected | 55.00 | 47.70 | 73.61 | 0.18 |
| 8 | 24 | Skin | Infected | 107.90 | 91.60 | 73.40 | 0.20 |
| 9 | 24 | Skin | Infected | 56.63 | 49.14 | 73.31 | 0.25 |
| 10 | 24 | Skin | Infected | 69.40 | 58.29 | 75.44 | 0.19 |
| 11 | 24 | Skin | Infected | 97.38 | 82.44 | 72.24 | 0.18 |
| 12 | 24 | Skin | Infected | 53.82 | 46.92 | 75.13 | 0.56 |
| 13 | 24 | Skin | Infected | 27.87 | 24.00 | 74.63 | 0.27 |
| 14 | 24 | Skin | Infected | 83.88 | 73.75 | 74.85 | 0.45 |
| 15 | 24 | Skin | Infected | 27.47 | 24.36 | 73.01 | 0.37 |
| 16 | 24 | Skin | Control | 29.82 | 26.31 | 72.21 | 0.06 |
| 17 | 24 | Skin | Control | 19.49 | 16.78 | 68.52 | 0.06 |
| 18 | 24 | Skin | Control | 25.83 | 22.92 | 72.44 | 0.05 |
| 19 | 24 | Skin | Control | 26.47 | 23.06 | 72.30 | 0.05 |
| 20 | 24 | Skin | Control | 25.14 | 21.95 | 71.70 | 0.06 |
| 21 | 24 | Skin | Control | 22.09 | 19.22 | 72.62 | 0.06 |
| 22 | 24 | Skin | Control | 24.47 | 21.51 | 73.42 | 0.07 |
| 23 | 24 | Skin | Control | 21.62 | 18.89 | 71.39 | 0.06 |
| 24 | 24 | Skin | Control | 29.19 | 25.68 | 70.18 | 0.05 |
| 25 | 24 | Skin | Control | 26.09 | 22.62 | 75.46 | 0.04 |
| 31 | 96 | Skin | Resistant | 27.73 | 24.35 | 72.16 | 0.04 |
| 32 | 96 | Skin | Resistant | 24.86 | 21.52 | 69.56 | 0.04 |
| 33 | 96 | Skin | Resistant | 38.30 | 33.21 | 68.28 | 0.05 |
| 35 | 96 | Skin | Resistant | 24.33 | 21.50 | 71.61 | 0.06 |
| 36 | 96 | Skin | Resistant | 26.68 | 23.52 | 68.68 | 0.05 |
| 37 | 96 | Skin | Resistant | 32.65 | 28.13 | 69.13 | 0.05 |
| 38 | 96 | Skin | Resistant | 23.91 | 20.64 | 58.65 | 0.06 |
| 39 | 96 | Skin | Resistant | 29.83 | 26.33 | 66.59 | 0.05 |
| 41 | 96 | Skin | Resistant | 27.47 | 22.18 | 71.01 | 0.06 |
| 42 | 96 | Skin | Resistant | 33.06 | 28.69 | 71.06 | 0.06 |
| 43 | 96 | Skin | Resistant | 28.96 | 25.06 | 70.41 | 0.06 |
| 44 | 96 | Skin | Resistant | 22.36 | 19.47 | 69.93 | 0.06 |
| 45 | 96 | Skin | Control | 26.62 | 19.43 | 74.74 | 0.06 |
| 46 | 96 | Skin | Control | 28.08 | 24.97 | 71.49 | 0.06 |
| 47 | 96 | Skin | Control | 20.88 | 16.62 | 70.30 | 0.04 |
| 48 | 96 | Skin | Control | 32.74 | 28.33 | 68.53 | 0.06 |
| 49 | 96 | Skin | Control | 35.34 | 31.52 | 71.05 | 0.05 |
| 50 | 96 | Skin | Control | 25.97 | 23.12 | 71.03 | 0.05 |
| 51 | 96 | Skin | Control | 19.73 | 13.76 | 69.39 | 0.06 |
| 53 | 96 | Skin | Control | 23.58 | 20.60 | 74.23 | 0.06 |
| 54 | 96 | Skin | Control | 31.75 | 27.35 | 69.89 | 0.06 |
| C2 | NA | T.maritimum | in culture | 32.44 | 26.57 | 73.17 | NA |
| C3 | NA | T.maritimum | in culture | 23.99 | 19.41 | 71.46 | NA |
| C4 | NA | T.maritimum | in culture | 28.02 | 22.88 | 66.10 | NA |
| C5 | NA | T.maritimum | in culture | 59.94 | 49.39 | 78.33 | NA |

**Table S 2: Individual sequencing and mapping statistics for MiSeq Illumina reads.** Hpi = Hours post-infection. PE = Paired-end.

| Alias | Condition | HPI | PE raw reads | PE filtered reads |
|---|---|---|---|---|
| T1_C1_1 | Control | 24 | 181621 | 145781 |
| T1_C1_2 | Control | 24 | 154374 | 130919 |
| T1_C1_3 | Control | 24 | 225514 | 190868 |
| T1_C1_4 | Control | 24 | 277890 | 225601 |
| T1_C1_5 | Control | 24 | 274337 | 231483 |
| T1_C2_1 | Control | 24 | 186787 | 143215 |
| T1_C2_2 | Control | 24 | 284429 | 233383 |
| T1_C2_3 | Control | 24 | 245827 | 201420 |
| T1_C2_4 | Control | 24 | 241375 | 198678 |
| T1_C2_5 | Control | 24 | 222868 | 182854 |
| T1_R1_F1 | Infected | 24 | 280961 | 220693 |
| T1_R1_F2 | Infected | 24 | 262423 | 207350 |
| T1_R1_F3 | Infected | 24 | 238551 | 194657 |
| T1_R1_F4 | Infected | 24 | 274968 | 208067 |
| T1_R1_F5 | Infected | 24 | 167319 | 141917 |
| T1_R2_F1 | Infected | 24 | 231367 | 196040 |
| T1_R2_F2 | Infected | 24 | 248824 | 199296 |
| T1_R2_F3 | Infected | 24 | 245764 | 198389 |
| T1_R2_F4 | Infected | 24 | 221779 | 181173 |
| T1_R2_F5 | Infected | 24 | 229554 | 196807 |
| T1_R3_F1 | Infected | 24 | 244098 | 206063 |
| T1_R3_F2 | Infected | 24 | 261797 | 221102 |
| T1_R3_F3 | Infected | 24 | 246235 | 205492 |
| T1_R3_F4 | Infected | 24 | 257414 | 214177 |
| T1_R3_F5 | Infected | 24 | 259603 | 216547 |
| T2_C1_1 | Control | 96 | 217474 | 175879 |
| T2_C1_2 | Control | 96 | 164151 | 132325 |
| T2_C1_3 | Control | 96 | 158113 | 129201 |
| T2_C1_4 | Control | 96 | 270518 | 223414 |
| T2_C1_5 | Control | 96 | 259156 | 219802 |
| T2_C2_1 | Control | 96 | 258307 | 218824 |
| T2_C2_3 | Control | 96 | 257713 | 210506 |
| T2_C2_4 | Control | 96 | 246841 | 209234 |
| T2_C2_5 | Control | 96 | 231808 | 193212 |
| T2_R1_F1 | Infected | 96 | 125049 | 105461 |
| T2_R1_F2 | Infected | 96 | 215821 | 188300 |
| T2_R1_F3 | Infected | 96 | 235308 | 197565 |
| T2_R1_F5 | Infected | 96 | 228482 | 190246 |
| T2_R2_F1 | Infected | 96 | 223099 | 187224 |
| T2_R2_F2 | Infected | 96 | 234896 | 193151 |
| T2_R2_F3 | Infected | 96 | 218448 | 180174 |
| T2_R2_F4 | Infected | 96 | 202990 | 162392 |
| T2_R3_F1 | Infected | 96 | 241555 | 204046 |
| T2_R3_F2 | Infected | 96 | 245816 | 207677 |
| T2_R3_F3 | Infected | 96 | 203071 | 165781 |
| T2_R3_F4 | Infected | 96 | 269876 | 204224 |
| T2_R3_F5 | Infected | 96 | 190538 | 163357 |

**Table S 3:** ***P. orbicularis*** **transcriptome statistics.**

| Transcriptome statistics | |
|---|---|
| Raw number of contigs | 211,840 |
| Total number of filtered contigs | 40,130 |
| Percent GC | 47.61 |
| Contigs N50 (bp) | 2,664 |
| Total assembled bases | 67,152,264 |
| Median contig length (bp) | 1,151 |
| Average contig length (bp) | 1,673.37 |
| **Annotation** | |
| Contig with match against nt (e-value 10-4) | 34,969 (87.14%) |
| Contig with match against Uniprot-Swissprot (e-value 10-3) | 27,801 (69.28%) |
| Contig with GO identifier annotation | 26,630 (66.36%) |
| Percent BUSCO completedness (metazoa_od10) | 93.6 |

**Table S 4: List of Fish and** ***T. maritimum*** **DEGs and GO terms enrichment**

Excel file: LeLuyer_etal_microbiome.TabS4.xls
